# The Duality of Adiponectin and the Role of Sex in Atherosclerosis

**DOI:** 10.1101/2023.05.23.541764

**Authors:** Abigail E. Cullen, Ann Marie Centner, Riley Deitado, Vladimir Ukhanov, Ahmed Ismaeel, Panagiotis Koutakis, Judy Delp, Gloria Salazar

## Abstract

Adiponectin, a hormone highly abundant in circulation, has many beneficial effects in atherosclerosis; however, gene deficiency of this hormone or its receptor have shown detrimental effects on plaque burden in mice. Our objective was to understand the role of sex and aging in the effects of adiponectin deficiency on plaque content, inflammation, and the mechanisms regulating the phenotype of *adipoq^-/-^* vascular smooth muscle cells (VSMCs). Even a 50% reduction in the expression of adiponectin led to a plaque reduction in males and an increase in females, compared with *apoe^-/-^*controls. Plaque reduction may be attributable to chemokines upregulated in males and downregulated in females. Changes in plaque were not attributed to changes in cholesterol or cardiovascular disease (CVD) markers. In old mice, both genotypes and sexes accumulated more plaque than *apoe^-/-^*. RNA sequencing of VSMCs from male mice in vitro uncovered a critical role for adiponectin in AKT signaling, regulation of the extracellular matrix, and TGF-β signaling. Upregulation of AKT activity mediated proliferation and migration of *adipoq^-/-^*cells. Activation of AMPK with metformin or AdipoRon reduced AKT-dependent proliferation and migration of *adipoq^-/-^* cells but did not improve the expression of contractile genes. Anti-atherogenic mechanisms targeted the ECM in *adipoq^-/-^* cells, downregulating MMP2 and 9 and upregulating decorin.

Our study uncovered sex and age-dependent effects of adiponectin deficiency in atherosclerosis.

## Introduction

Atherosclerosis, the major cause of cardiovascular diseases (CVDs), is exacerbated by risk factors including aging, obesity, and dietary choices. An additional risk factor arising from obesity and aging is the reduction of adiponectin (1, 2). Adiponectin, a hormone secreted in high levels by adipocytes, has been shown to positively alter the phenotype of vascular smooth muscle cells (VSMCs) (3); thus, the decreased circulating levels of adiponectin in disease states puts the body at heightened risk for other CVDs (4). Although the correlation between hypoadiponectinemia and atherosclerosis has been shown, specific mechanisms behind its role in mitigating of plaque formation have yet to be elucidated (5, 6).

Adiponectin production and secretion has historically been associated with adipose tissue; however, a multitude of tissues and cells, including skeletal and VSMCs, placental tissue, and cardiac cells, can also secrete the adipokine. Still, adipose tissue is the major source of adiponectin secretion (1, 3). Adiponectin binds to several receptors, including Adiponectin R1 and R2 (AR1 and AR2, respectively) and T-cadherin. AR1 or AR2 play a major role in VSMC’s contractile phenotype, while T-cadherin plays a more important role in angiogenesis and revascularization (7). Adiponectin mediates its effects on the metabolic state of the cell through activation of the AMP-activated protein kinase (AMPK), mitogen activated protein kinase (MAPK), and protein kinase B (AKT) pathways (8, 9). In the vascular system, adiponectin acts in a paracrine manner inducing the expression of contractile proteins leading to the differentiation of VSMCs through AMPK activation (3, 10). Thus, adiponectin promotes a contractile/anti-atherogenic phenotype in VSMCs. However, the role of adiponectin in atherosclerosis is unclear, as treatment with adiponectin reduces atherosclerosis in male apolipoprotein E (*apoe)^-/-^* mice (11), while lack of the adiponectin gene in low-density lipoprotein receptor (LDLR) *ldlr^-/-^* or *apoe^-/-^* mice showed no effect on plaque(12). In fact, plaque trended downwards in male *apoe*^-/-^*adipoq*^-/-^ mice. Both sexes were included for the *ldlr^-/-^*, but not for the *apoe^-/-^* background. Thus, the role of sex and the specific sites of plaque accumulation in the aortic tree in response to adiponectin deficiency needs further elucidation.

In this study, we used male and female *apoe^-/-^adipoq^-/-^*mice and uncovered sex– and age– dependent effects of adiponectin on plaque. Less plaque was seen in the arch and descending aorta of young *apoe^-/-^adipoq^-/-^* males treated with high fat diet (HFD) for 5 weeks, compared with *apoe^-/-^* controls. Interestingly, however, the opposite was observed in female mice. The protective effect of adiponectin deficiency in males was lost during aging, as 1-year-old males and females, fed a normal chow diet, had more plaque, compared with *apoe^-/-^* controls. RNA sequencing analysis of VSMCs isolated from the aorta of *adipoq^-/-^* males revealed a prominent role of adiponectin in the PI3K/AKT pathway, which was upregulated by adiponectin deficiency, inducing VSMC proliferation and migration. Our data show new insights into how adiponectin can regulate VSMC phenotype through different mechanisms. We show that adiponectin plays a role in the regulation of the extracellular matrix (ECM) through the regulation of matrix metalloproteinases (MMPs) and ECM components such as collagen and transforming growth factor-β (TGF-β). Further, unlike other studies, we have shown that there are sex–dependent differences in plaque as well as circulating pro-inflammatory molecules that warrant further investigation. We also show for the first time that even a 50% reduction in adiponectin, achieved using the *apoe^-/-^adipoq^+/-^* mouse, has significant impact on the development of atherosclerosis. Further, our study suggests that females with reduced adiponectin levels may be at a higher risk of developing atherosclerosis.

## Results

### Plaque is reduced in adiponectin deficient males but increased in females

To test the role of adiponectin and sex in atherosclerosis, male and female *apoe^-/-^ adipoq^+/+^, apoe*^-/-^*adipoq*^+/-^, and *apoe*^-/-^*adipoq*^-/-^ were fed HFD for 5 weeks ad libitum. In contrast to previous reports (12), en face analysis (**Fig. 1A-C)** of the aortas of male *apoe^-/-^adipoq^+/-^*and *apoe^-/-^adipoq*^-/-^ mice indicated a significant reduction in plaque in the arch (p=0.037 and 0.028, respectively) and descending aorta (p=6.2E-5 and 0.004, respectively), compared with *apoe^-/-^* controls. In contrast, females had more plaque in the arch for both genotypes (p= 0.013 and 0.014, respectively), and more plaque in the descending aorta of *apoe^-/-^adipoq^+/-^* mice (p=0.03). A trend to higher plaque was also seen in the descending aorta of *apoe^-/-^adipoq*^-/-^ female mice. Due to the beneficial impact of estrogen on the cardiovascular system (13), we analyzed estradiol levels using Luminex technology (**Fig. 1D**). We used plasma from the *apoe^-/-^*control and *apoe*^-/-^ *adipoq*^-/-^ mice for this analysis. Consistent with the reduction of plaque, estradiol was elevated in males lacking adiponectin compared with *apoe^-/-^* controls (p=0.04). A trend toward higher levels of estrogen was seen for *apoe^-/-^adipoq*^-/-^ female mice. The sex-dependent effects in plaque did not correlate with changes in HFD intake, water consumption, or body weight (BW) in males (**Fig. S1A-C)** or females **(Fig. S1D-F).** In fact, in week 5, the heterozygous male group ate significantly more than the *apoe^-/-^* and *apoe^-/-^adipoq*^-/-^ groups (p=0.017 and p=0.02, respectively), and in week 2, the *apoe^-/-^adipoq*^-/-^ drank more water than the *apoe^-/-^* and *apoe^-/-^adipoq*^+/-^ groups (p=0.006 and 0.04, respectively). There were no significant differences in food and water consumption or body weight for the female groups fed HFD.

**Figure 1.**
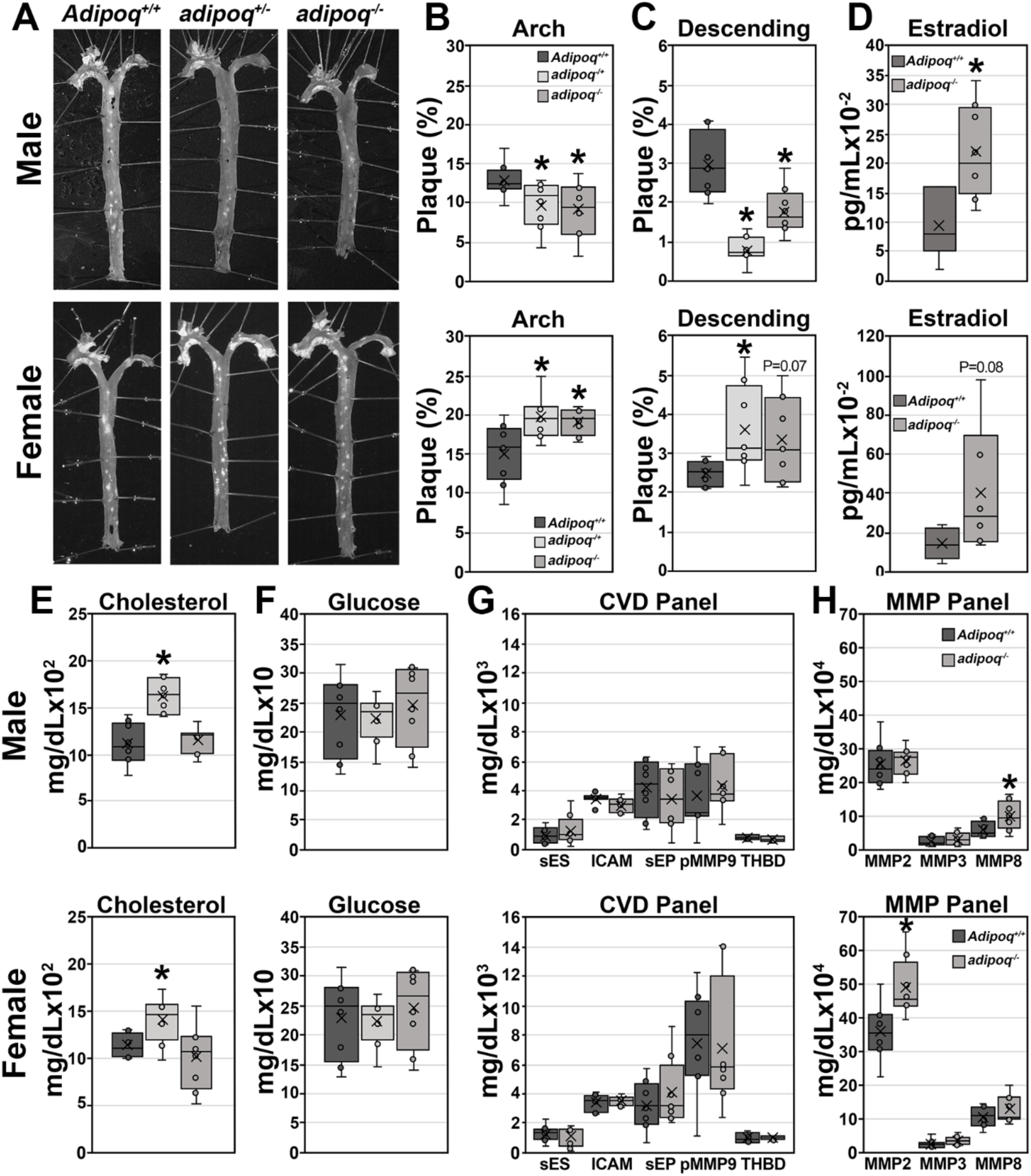
Loss of adiponectin reduces plaque accumulation in males, but not females. En face analysis of plaque accumulation in male and female *apoe^-/-^adipoq^+/+^, apoe^-/-^adipoq^+/-^*, and *apoe^-/-^adipoq^-/-^*after 5 weeks of HFD treatment (n=8 per group for males and 7-9 per group for females) (A). Plaque was quantified in the arch (B) and descending aorta (C). Serum from males and females *apoe^-/-^adipoq^+/+^* and *apoe^-/-^adipoq^-/-^* was tested for estradiol (D), cholesterol (CHO) (E), glucose (F), CVD markers (G) and MMPs (H). * denotes p < 0.05, compared with *apoe^-/-^* control.

The sex-dependent effect of adiponectin in plaque was also not correlated with circulating cholesterol (CHO) or glucose. Surprisingly, CHO was elevated in both sexes in heterozygotes, (**Fig. 1E**) and glucose did not differ between groups (**Fig. 1F**). These results indicate that the loss of adiponectin did not promote atherosclerosis through any of these prominent pro-atherogenic pathways. To get insights into possible mechanisms, we analyzed CVD markers. No differences were seen between *apoe^-/-^* and *apoe^-/-^adipoq*^-/-.^ in circulating levels of soluble E and P selectins (sES, sPS), soluble intercellular adhesion molecule 1 (sICAM-1), pro-matrix metalloproteinase (MMP) 9, or thrombomodulin (THBD) in males or females (**Fig. 1G**). For the MMP panel, *apoe^-/-^adipoq*^-/-^ males had higher levels of MMP8 (p=0.03), while females had elevated MMP2 (p<0.01) and a trend towards higher MMP3 (p=0.058), compared with *apoe^-/-^* controls (**Fig. 1H**).

Overall, the lack of adiponectin had a protective effect in HFD-induced plaque in males, but not in female mice. However, the adaptation of young males fed HFD was not seen in 1-year-old animals maintained on standard chow diet, as both males and females of both genotypes exhibited higher plaque content in the arch, compared with *apoe^-/-^* controls (**Fig. S2A and B**). Only aged female *apoe^-/-^adipoq^+/-^* mice showed more plaque in the descending aorta (**Fig. S2A and C**).

### Adiponectin has sex-dependent effects on inflammation

To determine if the changes seen in plaque by HFD in young mice were associated with changes in inflammation, we used Luminex technology to assess the expression of pro-inflammatory cytokines and chemokines, focusing only on double knock-outs (KO) (**Fig. 2)**. There were no significant changes between sex or genotypes for Interleukin (IL)-2 or –9, eotaxin/CC motif chemokine (CCL) 11, monokine-induced by gamma interferon (MIG)/ C-X-C motif chemokine (CXCL) 9, macrophage inflammatory marker (MIP2)/CXCL2, or regulated on activation, normal T cell expressed and secreted (Rantes)/CCL5 (**Fig. 2B, F, I, O, Q, and R**).

**Figure 2.**
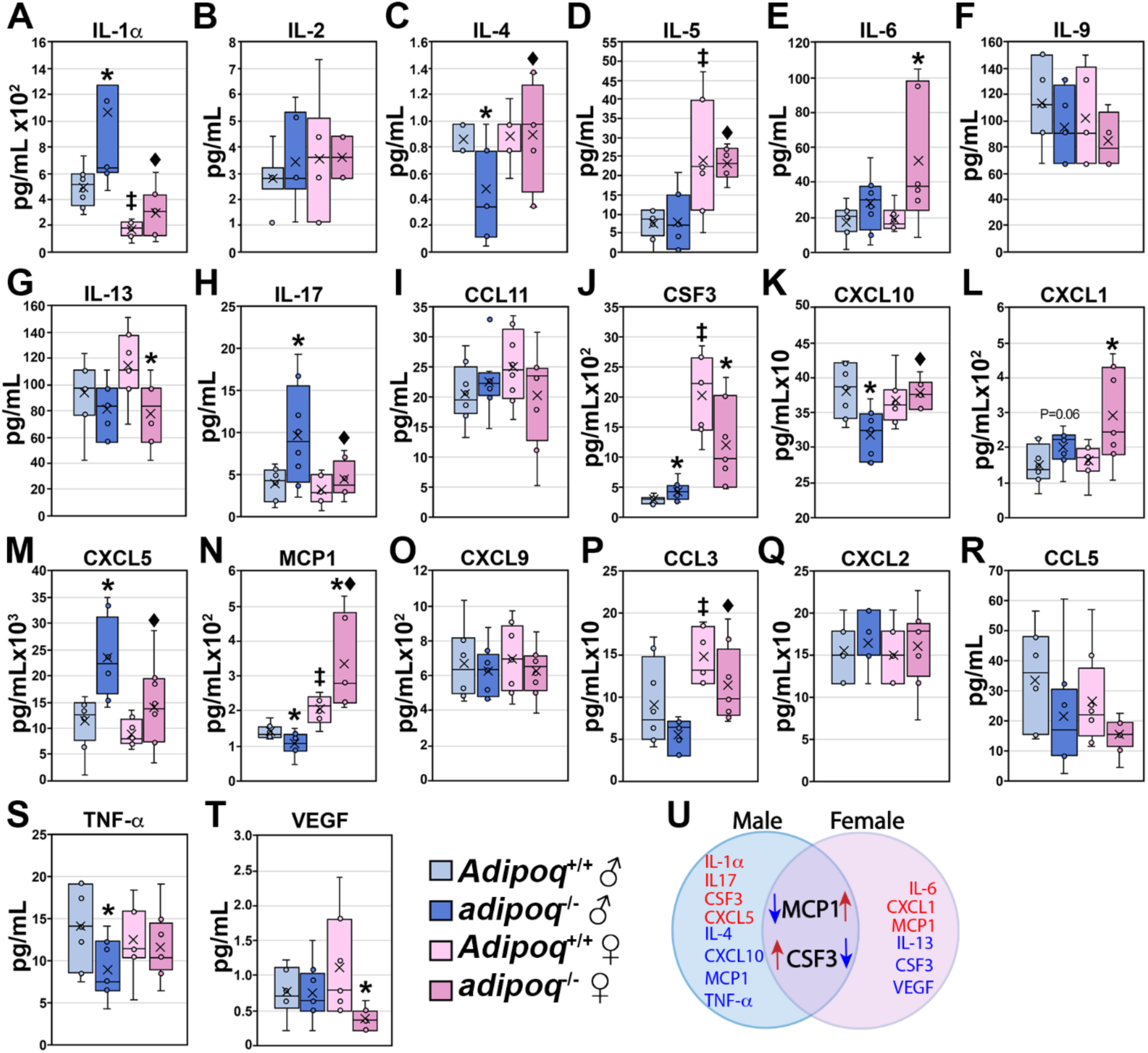
Inflammatory markers and chemokines are influenced by adiponectin and sex. Serum samples from male and female *apoe^-/-^adipoq^+/+^* and *apoe^-/-^adipoq^-/-^*animals treated for 5 weeks with HFD were tested for the expression of inflammatory markers and chemokines using the Milliplex mouse cytokine/chemokine panel 1 (n=5-8 per group) (A-T). Overall changes are summarized in U. * denotes p < 0.05 for genotype-dependent changes within sex, ‡ denotes p < 0.05 for sex dependent changes between *Adipoq^+/+^* animals, and diamond denotes p < 0.05 for sex dependent differences between *adipoq^-/-^*animals.

Changes seen in male by adiponectin deficiency included upregulation of IL-1α, IL-17, granulocyte colony-stimulating factor (G-CSF)/CSF3, CXCL5 and a trend for CXCL1 (**Fig. 2A, H, J, M and L**). A downregulation in expression was seen for IL-4, CXCL10/interferon gamma-induced protein 10 (IP-10), monocyte chemoattractant protein 1 (MCP1) and tumor necrosis α (TNF-α) **(Fig. 2C, K, N and S**). In females, the lack of adiponectin upregulated IL-6, keratinocyte chemoattractant (KC/CXCL1) and MCP1 (**Fig. 2E, L, and N**) and downregulated IL-13, CSF3, and vascular endothelial growth factor (VEGF) (**Fig. 2G, J, and T**). Compared with males, females in both groups had reduced IL-1α and elevated IL-5, CSF3, MCP1 and macrophage inflammatory protein-1α (MIP1α/CCL3) (**Fig. 2A, D, J, N, and P**). Altogether, we observed genotype and sex-dependent differences in inflammatory markers (**Fig. 2U**). The downregulation in IL-4, CXCL10, MCP1 and TNF-α and estradiol upregulation may mediate the adaptation of young adiponectin deficient males to HFD-induced atherosclerosis.

### Adiponectin deficiency upregulates the AKT signaling pathway and reduces VSMC differentiation

To get insights into the possible protective mechanisms seen in males, we isolated VSMCs from wild type (WT) and *adipoq^-/-^* male mice and performed RNA sequencing analysis. The results yielded 15,986 genes, of which 1,054 were upregulated and 1,187 were downregulated in *adipoq^-/-^*cells, compared with WT (**Fig. S3A**). Differentially expressed genes (DEGs) were labeled as up or down if the log2 of the fold change (*adipoq^-/-^*/*Adipoq^+/+^*) was higher or lower than 0.58, respectively. The most upregulated gene was the cell adhesion molecule contactin associated protein 2 (Cntnap2, ∼600-fold), while the most downregulated gene was the complement component 4 (C4b, ∼270-fold) followed by fibromodulin (Fmod, ∼80-fold) and Apoe (∼70-fold).

Gene ontology analysis of the DEGs showed many genes were involved in cellular processes, human diseases, and metabolism (**Fig. S3B**). For example, 387 genes were associated with signal transduction, while 236 were involved in cancer. Further analysis of the most enriched pathways (**Fig. S3C**) revealed prominent changes in the phosphoinositide-3-kinase (PI3K)-AKT signaling pathway with ∼40 genes upregulated and ∼50 downregulated. Some of the most enriched (Kyoto encyclopedia of genes and genomes) KEGG pathways (**Fig. 3A**) affected also included the Ras and MAPK cascades, cell adhesion, focal adhesion, ECM-receptor interaction, pathways in cancer, axon guidance, and complement and coagulation cascades. Network analysis (**Fig. S3D**) of these pathways showed close interactions among the PI3K-AKT, pathways in cancer, Rap1 signaling and the ECM-receptor interaction networks. Many genes in the cancer network overlap with the PI3K/AKT, as both pathways regulate cell growth. Further analysis of the ECM-receptor interaction, MAPK signaling, and PI3K/AKT pathways indicate many alterations in the absence of adiponectin. The most significant changes are indicated by arrows on the heatmaps shown in **Fig. 3B**.

**Figure 3.**
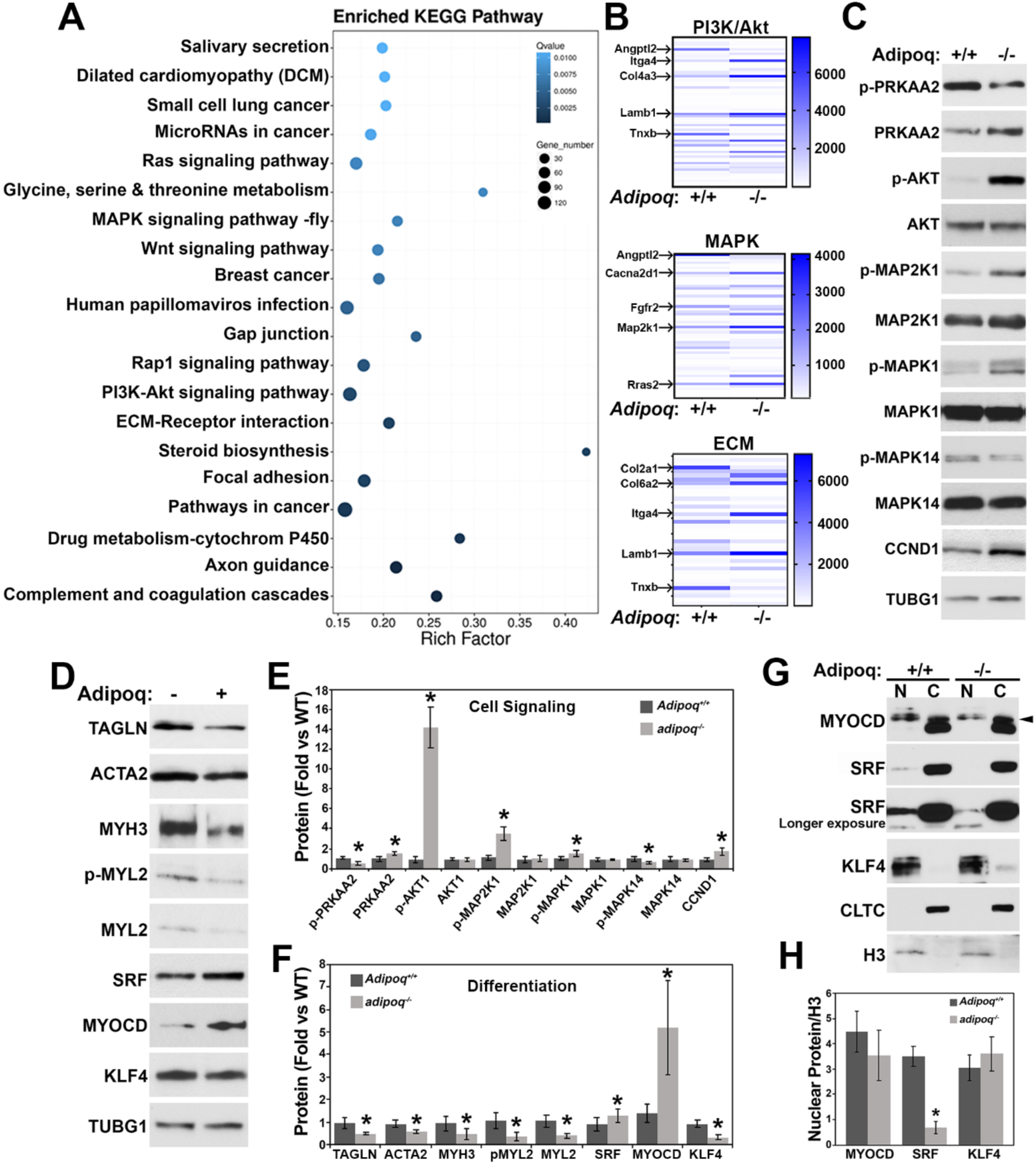
Vascular smooth muscle cell phenotype is altered by loss of adiponectin. RNA sequence data results showed the most enriched KEGG pathways in the *adipoq^-/-^*cells (A) and heat maps indicate the specific genes altered in the PI3K/AKT, MAPK, and ECM pathways (B). Confluent *Adipoq^+/+^* and *adipo^-/-^*cells were starved in 0.2% FBS for 24 h and total extracts (30-50 μg) tested for signaling cascade (C) and differentiation markers (D). Respective quantification graphs are shown in E and F. Nuclear and cytosolic fractions (20 μg) were tested for MYOCD, SRF and KLF4 (G and H) * denotes p < 0.05 compared to *Adipoq^+/+^*.

Next, we focused on AKT and MAPK signaling pathways that are known to interact with AMPK, the major downstream effector of the adiponectin receptor and compared the expression from the RNA sequencing (**Fig. S4A**) with the protein levels (**Fig. 3C and E**). Phosphorylation of the AMPK catalytic subunit (p-PRKAA2) was reduced by 50%, while total levels were upregulated in *adipoq^-/-^* cells. Consistent with reduced AMPK activity, AKT phosphorylation was strongly upregulated ∼14-fold, which could be mediated by the upregulation of Akt1 mRNA. *adipoq^-/-^* cells also expressed higher levels of phosphorylated mitogen-activated protein kinase kinase 1 (MAP2K1/MEK) and its downstream kinase MAPK1/ERK. Because AKT and MAPK1 negatively impact AMPK activity (14, 15), our data suggest that a negative loop mediated by these kinases may contribute to AMPK reduced activity. In addition to the upregulation of proliferative signaling pathways, protein, but not RNA expression of the cell cycle progression kinase cyclin D1 (CCND1) was also increased by the lack of adiponectin. In addition to the AMPK pathway, activation of AR1 and AR1 receptors, also stimulates p38MAPK (MAPK14)(16). Consistent with the lack of adiponectin MAPK14 was phosphorylation was significantly reduced in the KO cells.

VSMCs have been shown to secrete adiponectin, which promoted a contractile phenotype in VSMCs (3). In line with this report, in *adipoq^-/-^* VSMCs RNA sequence (**Fig. S4B**) and western blot data (**Fig. 3D and F**) show reduced expression of contractile genes including transgelin (TAGLN/SM22), smooth muscle actin (ACTA2), smooth muscle myosin heavy chain (MYH), and myosin light chain (MYL).

Contractile genes are regulated by the transcriptional coactivator myocardin (MYOCD) and transcription factor serum response factor (SRF) (17). Negative regulators of MYOCD, such as Krüppel like factor 4 (KLF4) prevent the formation of the MYOCD/SRF complex, reducing the expression of differentiation genes. Interestingly, while the differentiation markers tested were significantly reduced in *adipoq*^-/-^ cells, MYOCD and SRF were upregulated at the RNA (**Fig. S4B)** and protein levels (**Fig. 3D and F)**. RNA and protein levels for KLF4 were increased and reduced, respectively in the *adipoq*^-/-^ cells. (**Fig. S3)**. Next, we tested whether altered subcellular localization of these genes may explain the reduced expression of differentiation markers. In fact, *adipoq*^-/-^ cells showed a robust reduction in nuclear SRF and no differences in the level of MYOCD or KLF4 in the nucleus (**Fig. 3G and H**). Clathrin heavy chain (CLTC) and histone H3 (H3) were used as cytosolic and nuclear markers, respectively.

### Loss of adiponectin promotes a synthetic phenotype altering ECM composition

Increased cell signaling of the AKT and MAPK pathways, and reduced expression of contractile genes induced a switch toward a synthetic phenotype. The logarithmic phase of growth of VSMCs was increased in the *adipoq^-/-^* cells (**Fig. 4A**), reaching a higher cell density at days 5 and 6, compared with *Adipoq^+/+^* cells. Since both cell types reached confluency, these data suggest that *adipoq^-/-^* cells may be smaller in size. To discern if migration was also increased in parallel to proliferation, in the *adipoq^-/-^* cells, we measured scratch wound closure at 12 and 24 h (**Fig. 4B**). At both time points there was a significant increase in the % closure of the KO cells, potentially signifying differences in ECM regulation and motility regulators, like Rac2, as suggested by RNA sequence data (**Fig. S4C**). The switch toward the synthetic phenotype in *adipoq^-/-^*cells was associated with impaired oxidative phosphorylation (OXPHOS), specifically in complexes I and IV (**Fig. 4C**) and by upregulation of RNA expression of glycolytic enzymes (**Fig. S4D**). We confirmed the upregulation of protein expression of glycolytic enzymes including enolase 3 (ENO3), phosphofructokinase liver type (PFKL), a Pfk1 isoform, and phosphoglycerate mutase 1 (PGAM1) (**Fig. 4D and E**). AKT phosphorylates PFK isoforms, increasing its protein stability and proliferation rate in glioblastoma cells (18), suggesting that increased AKT activity may induce PFK expression and upregulate glycolysis in VSMCs. Similarly, lactate dehydrogenase (LDH), the enzyme that converts pyruvate into lactate, was also upregulated in *adipoq^-/-^*cells, suggesting that the NAD produced in this reaction may further drive glycolysis. Consistent with the altered function of complexes I and IV of the ETC, *adipoq*^-/-^ cells have higher levels of mitochondrial ROS (MitoROS), as well as superoxide (O_2_^.-^) and hydrogen peroxide (H_2_O_2_) (**Figure 4F**). Altogether these data suggest that *adipoq^-/-^* cells use glycolysis instead of OXPHOS to fuel their high rate of proliferation and migration.

**Figure 4.**
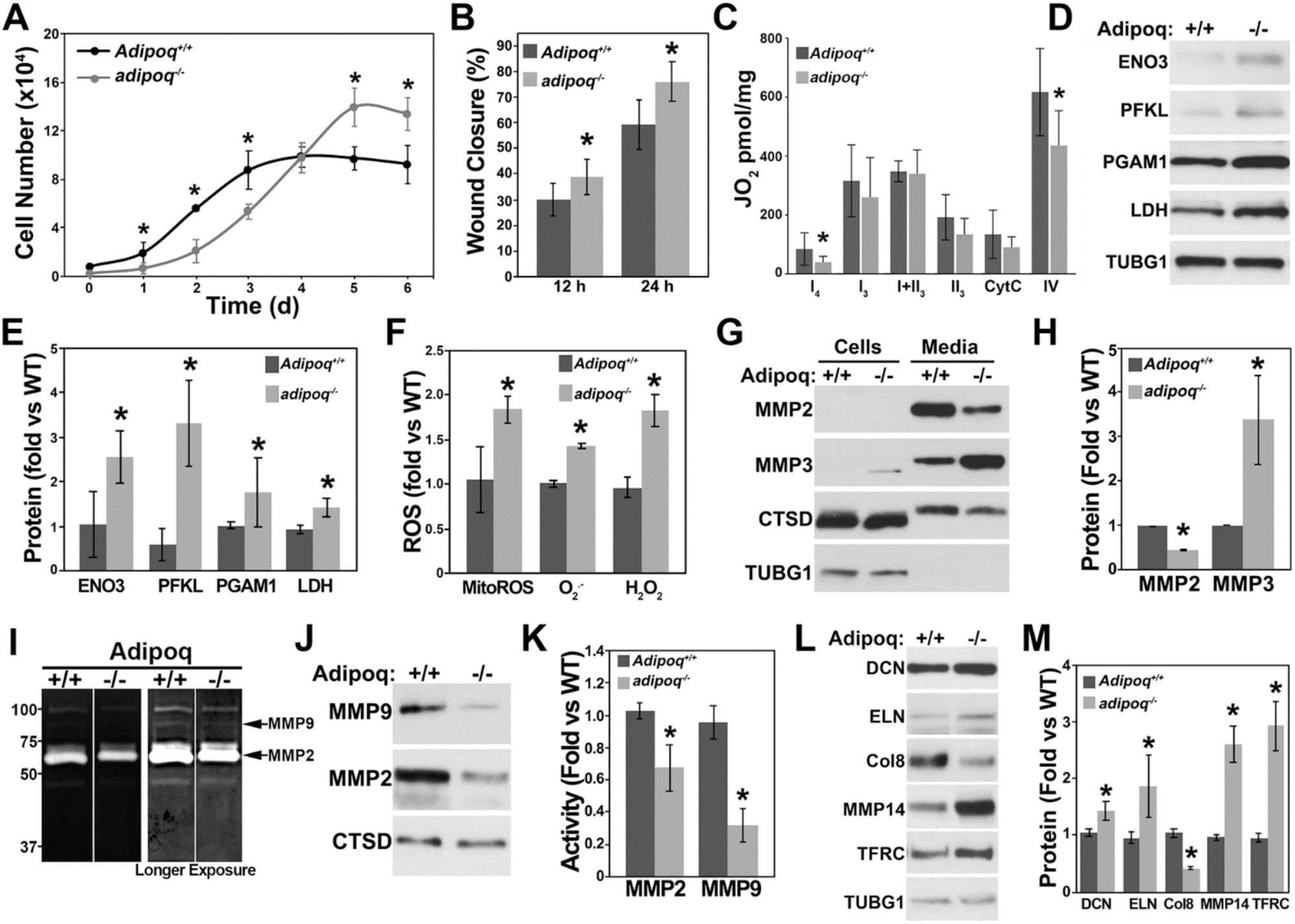
Loss of adiponectin alters metabolic activity, mitochondrial function, and ECM regulation. *Adipoq^+/+^* and *adipo^-/-^* cells were counted every day for 6 days to assess proliferation rate (n=3-4 wells per experiment) (A). Migration was assessed by scratch wound closure assay (B). Oxygen flux (JO2) was assessed by high-resolution respirometry using the Oroboros O2k Oxygraph. Complex I, state 2 (I_2_) respiration and state 3 respiration rates of Complex I (I_3_), combined Complex I and Complex II (I+II_3_), Complex II (II_3_), and Complex IV (IV_3_) were assessed (C). *Adipoq^+/+^* and *adipo^-/-^* cells were cultured to confluency then starved for 24 h in 0.2% FBS for western blot analysis of glycolytic enzymes (n=3-4) (D and E). *Adipoq^+/+^* and *adipo^-/-^* cells were starved for 24 h in FBS-free media for analysis of membrane bound and secreted MMPS (n=3-4) (G and H). The media was also used for gelatin zymography analysis of MMP activity (I). Western blot analysis of the same media was done to analyze expression of MMPs (J). Expression of ECM proteins and TFRC in total extracts of *Adipoq^+/+^* and *adipo^-/-^*cells (L and M). * denotes p < 0.05 compared to *Adipoq^+/+^*.

Synthetic VSMCs remodel the ECM by secreting MMPs and ECM components, stimulating migration (19). High expression of MMPs, in particular MMP2 and MMP9, degrade collagen and ELN, weakening the arterial wall contributing to plaque formation (20). However, at physiological/low levels MMPs can be beneficial because when the ECM is not degraded, arteries can begin to stiffen(21). To assess whether molecules involved in ECM remodeling were affected in the *adipoq*^-/-^ cells, the RNA sequence was searched for MMPs (**Fig. S4E**). Genes encoding MMPs 3, 19, 25, and 28 showed the strongest change relative to WT cells. MMP2 and MMP14 were expressed at the highest level based on RNA expression. Compared with WT cells, MMP2 was reduced by about 50% and MMP14 was not affected in *adipoq^-/-^* cells. In contrast, MMPs 3, 9, 15, and 23, 25 and 27 were upregulated. MMP9, however, was expressed at very low levels. Consistent with the RNA data, MMP2 was reduced while MMP3 was increased in the media collected from cultured *adipoq^-/-^*, compared with WT cells (**Fig. 4G and H**). Cathepsin D (CTSD) is secreted into the media and served as a loading control. Gelatinase activity measured by zymography showed bands of about 50 and 90 kDa, consistent with MMP2 and MMP9 molecular weights, that were reduced in *adipoq^-/-^* cells (**Fig. 4I**). Similar to MMP2, MMP9 was also reduced in the media of these cells (**Fig. 4J**), suggesting that reduced MMP2 and MMP9 activities may be mediated by lower protein levels. MMP9 degrades elastin (ELN) (22), while MMP2 and MMP3 degrade decorin (DCN). Further, collagen type VIII (Col8) has been shown to signal through β1 integrins to upregulate MMP2 and induce migration of smooth muscle cells (22). In line with this evidence, reduced MMP2 and MMP9 activities correlated with DCN and ELN upregulation at the RNA (**Fig. S4C**) and protein levels (**Fig. 4L and M**). Col8 RNA, on the other hand, was increased, while its protein level was reduced.

TGF-β is a major regulator of the ECM (23) and mediates VSMC differentiation (24). Activation of the TGF-β1 receptor stimulates pathways that initiate the production of ECM components such as collagen (25). The altered phenotype of *adipoq^-/-^* VSMCs suggest that TGF-β signaling could be impaired in these cells. Many regulators of the ECM and iron metabolism, such as the membrane type 1-matrix metalloproteinase (MT1-MMP/MMP14) and the transferrin receptor (TFRC) mediate TGF-β effects. MMP14 releases TGF-β from its binding to DCN in the ECM (26) and also releases TGF-β from latent transforming growth factor-β (LTBP) (27). DCN binds to TGF-β, preventing its binding to surface receptors, thus, acting as an antifibrotic molecule. TFRC, on the other hand, enables TGF-β –induced craniofacial morphogenesis (28) and TGF-β-induced renal fibrosis (27). Evaluation of the TGF-β pathway in the RNA Sequence (**Fig.S4F**) revealed upregulation of the Tfrc, transferrin (Trf), and Ltbp1, downregulation of Ltbp2 and Ltbp4 and no change in MMP14 genes (**Fig. S4E**) by adiponectin deficiency. However, at the protein level, both TFRC and MMP14 were upregulated (**Fig. 4L and M**).

Analysis of TGF-β-induced signaling revealed higher phosphorylation levels of AKT and suppressor of mothers against decapentaplegic (SMAD) 2 and higher expression of TFRC and MMP14 in *adiopq^-/-^* cells, compared with WT (**Fig. 5A, B and D-F**). Phosphorylation of MAPK1 was reduced in WT cells by TGF-β treatment, which correlated with upregulation of ACTA2. Surprisingly, TGF-β failed to reduce MAPK1 activity and upregulate ACTA2 in *adiopq^-/-^* cells (**Fig. 5A, C and G**).

**Figure 5.**
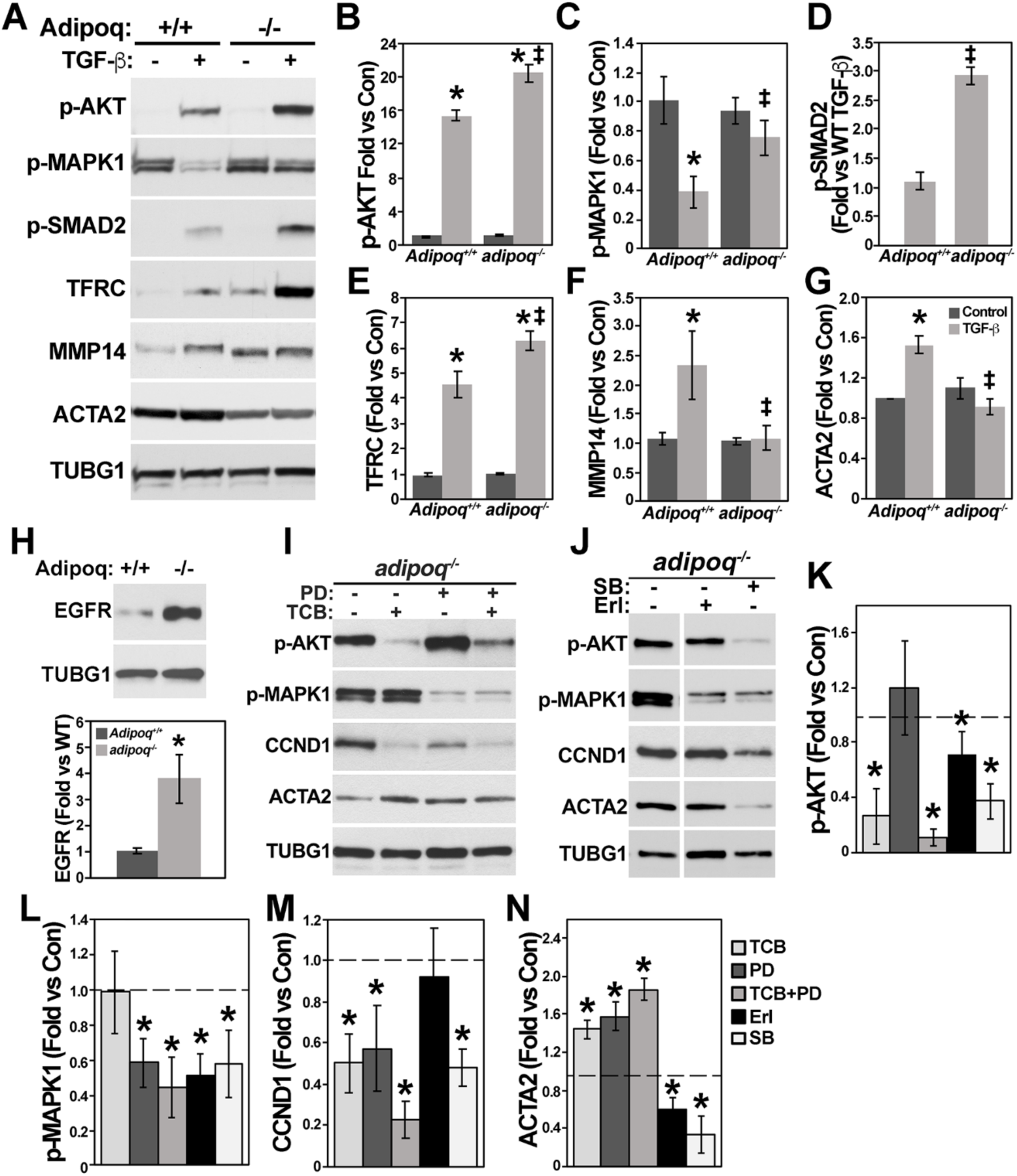
AKT and MAPK1 regulates the expression of differentiation and proliferation markers. *Adipoq^+/+^* and *adipoq ^-/-^* cells starved for 24 h with 0.2% FBS were treated with 10 ng/ml TGF-β for 24 h to test the expression of in cell signaling, ECM regulation, and differentiation markers (A-G) or to measure EGFR expression (H). Cells were treated with 10 μM TBC and PD-98059, 100 nM Erl or 5 μM SB-252334 for 24 h and total cell extracts used for western blot (I-N). In B-G * denotes p < 0.05 compared to *Adipoq^+/+^* or *adipo^-/-^* controls and ‡ between TGF-β-treated groups. In K-N * represent p<0.05 compared with *adipoq ^-/-^* control cells. (n=3-4 independent experiments).

We next investigated whether activation of AKT by TGF-β is mediated via transactivation of receptor tyrosine kinases like EGFR, which is also upregulated in *adipoq^-/-^* cells (**Fig. 5H**). Treatment with EGF showed similar responses for AKT and MAPK1 phosphorylation (**Fig. S5 A-C**) and similar levels of EGFR at the cell membrane in both genotypes (**Fig. S5D and E**), suggesting that upregulation of AKT activity is not caused by EGFR signaling. Caveolin 1 (CAV1) expression, on the other hand, was reduced in cell extracts and at the cell surface (**Figure S5D-F**). Interestingly, CAV1 inhibits SMAD2 dependent TGF-β signaling (29), suggesting that downregulation of CAV1 may promote the higher SMAD2 signaling observed in *adipoq^-/-^* cells. CTLC, which is required for EGFR internalization was unaffected by the lack of adiponectin.

These alterations suggest that the ECM surrounding *adipoq^-/-^*cells is structurally different from *Adipoq^+/+^* ECM, which in turn changes the rate of migration and proliferation, and differentiation of the *adipoq^-/-^* cells. Altogether these data suggest that DCN may serve as a reservoir for TGF-β, which can subsequently be released from the ECM by MMP14 and interact with the cell surface receptors to activate AKT and induce proliferation.

### AMPK inhibits AKT-dependent cell signaling, proliferation and migration but not differentiation of VSMCs

To determine the impact of AKT and MAPK signaling in the phenotypic switch, *adipoq^-/-^* cells were treated with TCB and PD. TCB and PD reduced CCND1 and elevated ACTA2, and their combination yielded the greatest effect on CCND1 and ACTA2 levels (**Fig. 5I and K-N**). These data suggest that upregulation of both AKT and the MAPK pathways promotes the synthetic phenotype of *adipoq^-/-^* cells. Inhibition of the EGFR by Erl showed a weaker effect in AKT and MAPK1 phosphorylation and failed to reduce CCND1 and increase ACTA2. Inhibition of TGF-β signaling by SB reduced all of these markers (**Fig. 5J-N**). Thus, inhibition of AKT and MAPK1 and activation of the TGF-β pathways are likely needed to maintain the contractile phenotype of VSMCs.

Next, we investigated whether activation of adiponectin signaling can downregulate AKT and MAPK1 pathways and reverse the phenotypic switch of the *adipoq*^-/-^ cells. We tested this pathway by activating adiponectin receptors with conditioned media from WT cells that are known to secrete adiponectin and with recombinant adiponectin. Conditioned media from *Adipoq^+/+^* and *adipoq*^-/-^ cells showed a similar secretory profile. Silver staining detected a band around 30 kDa that was present in *Adipoq^+/+^* cells and was missing in the *adipoq*^-/-^ sample (**Fig. 6A, arrow**). Western blot of the same samples confirmed that this band was in fact adiponectin (**Fig. 6A**). Treatment of *adipoq*^-/-^ cells with conditioned media from WT cells showed reductions in AKT and MAPK1 phosphorylation as well as CCND1 and no changes in ACTA2 (**Fig. 6B and D**). Similar effects on AKT, MAPK1 and ACTA2 were seen in *adipoq^-/-^* cells treated with recombinant adiponectin (**Fig. 6C and D**). CCND1, however, was not downregulated.

**Figure 6.**
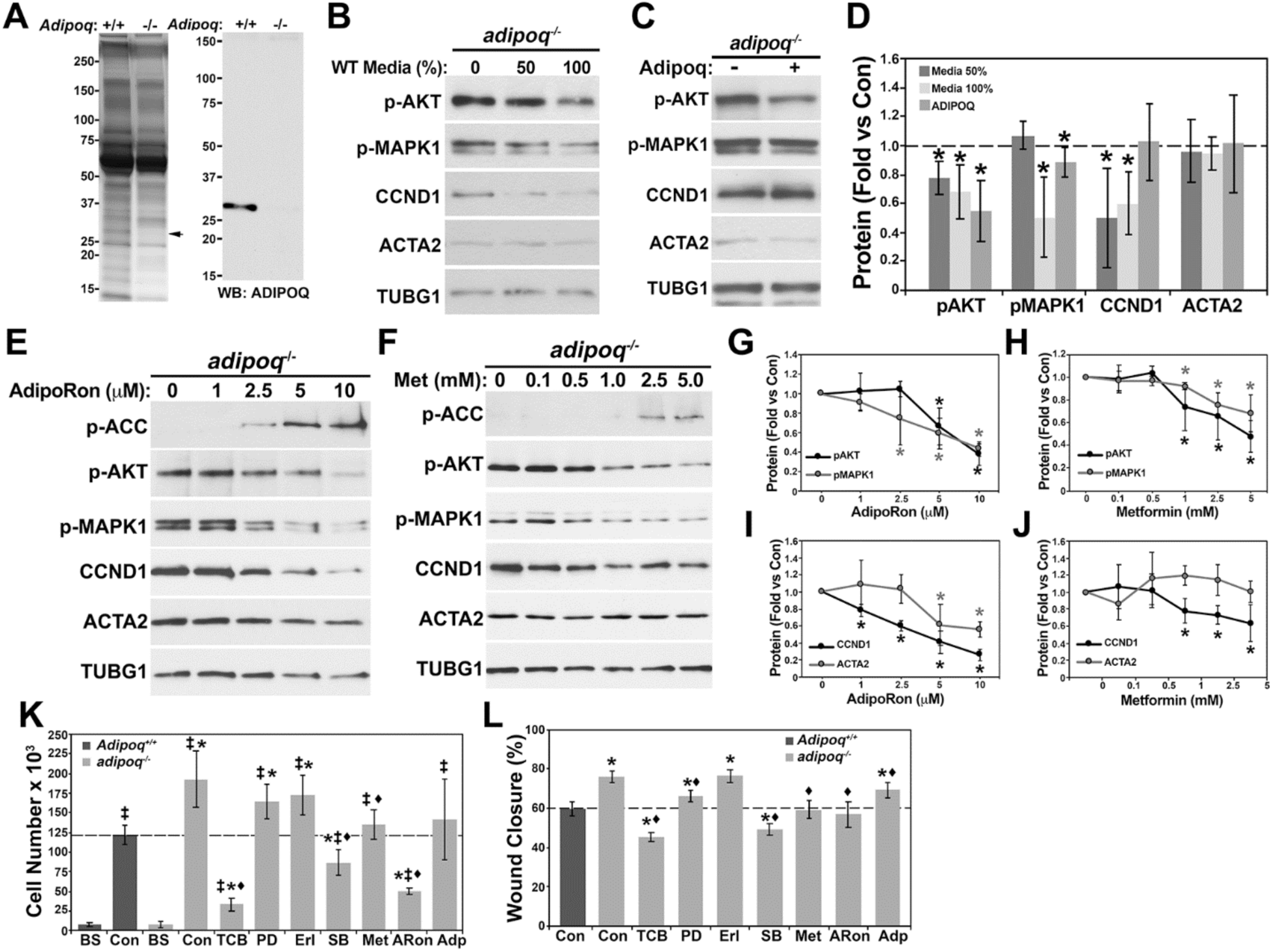
Restoration of adiponectin and AMPK activity reduces cell signaling but does not improve differentiation. Concentrated media from *Adipoq^+/+^* and *adipoq^-/-^* cells were separated in a 4-20% pre-cast Criterion gel and stained with silver or used for western blot to measure adiponectin (A). *adipoq^-/-^* cells were treated for 48 h with and without undiluted WT media (100%) or diluted 50% in plain media (B) or treated with 10 μg/mL recombinant adiponectin (C). Quantification for B and C is shown in D. *adipoq^-/-^* cells were treated for 72 h with AdipoRon (1-10 μM) or metformin (0.1-5 mM) (E-J). *adipoq^-/-^*cells were treated with 10 μM TBC, 10 μM PD-98059, 100 nM Erl, 5 μM SB-252334, 5 mM metformin, 10 μM AdipoRon or 10 μg/mL recombinant adiponectin for 4 days to measure proliferation (K) or for 1 d to measure migration (L). In K cells were seeded at low confluency, starved for 48 h and counted to establish basal (BS) cell numbers before treatment. For G-H * denotes p < 0.05 compared to *adipoq^-/-^* control. For K and L * represent p<0.05 compared with WT control, ‡ compared with BS for each genotype and ♦ compared with *adipoq^-/-^* control.

Adiponectin has been shown to activate T-cadherin and AR1/AR2 in both mouse and human cell lines (30, 31). To investigate whether activation of AR1 and AR2 mediates the effects of adiponectin in cell signaling, we used AdipoRon, an AR1 and AR2 agonist. AdipoRon increased the phosphorylation of the AMPK substrate acetyl-CoA carboxylase (ACC) and reduced AKT and MAPK1 phosphorylation and CCND1 in a concentration dependent manner, without affecting ACTA2 (**Fig. 6E, G and I**). Further, activation of AMPK with metformin mimicked the effects of AdipoRon (**Fig. 6F, H and J**). Altogether, these data suggest that adiponectin secreted by VSMCs activates AR1 and AR2, which mediates the downregulation of AKT and MAPK1 signaling and CCND1 in an AMPK-dependent manner. However, it is unknown, why the reduction in AKT and MAPK1 activities by AdipoRon and metformin were unable to upregulate ACTA2.

### AKT mediates the proliferation and migration of *adipoq^-/-^* cells

The basal rate of proliferation (**Fig. 4A**) was significantly higher in *adipoq**^-/-^*** cells compared to WT cells. Inhibition of AKT, MAPK1, and TGF-β and activation of AMPK (adiponectin, AdipoRon and metformin) reduced CCND1 expression, suggesting that these treatments should also reduce proliferation of *adipoq**^-/-^*** cells. WT and *adipoq^-/-^* cells were counted at basal (BS) and after 4 days of treatment with control media, media containing inhibitors of the AKT/MAPK1/TGF-β pathways or media containing activators of adiponectin signaling (**Fig. 6K**). The strongest reduction in proliferation was seen in cells treated with TCB, followed by AdipoRon and SB, all reaching cell numbers lower than WT cells. Metformin reduced proliferation to levels similar to WT cells. Inhibition of MAPK1 and EGFR, or treatment with adiponectin, led to reductions in cell number that did not reach significance. Similar to their effects on proliferation, TCB and SB treatment had the strongest effect on reducing migration of *adipoq*^-/-^ cells to levels lower than WT cells. Metformin and AdipoRon showed similar reductions in migration consistent with WT levels. Further, although PD and adiponectin reduced migration of *adipoq*^-/-^ cells, migration was still higher in comparison to WT cells. Inhibition of the EGFR by Erl showed no effect on migration. Altogether, these data suggest that upregulation of AKT signaling is the major contributor to the elevated rate of proliferation and migration of *adipoq^-/-^* cells.

### Loss of adiponectin dysregulates cellular signaling in males and females

To assess the significance of the molecular changes observed *in vitro*, we measured expression of proteins of interest in the aortas of male (n=8 per genotype) and female (n= 7 per genotype) *Adipoq^+/+^* and *adipoq^-/-^* mice (**Fig. 7**). Aortas of *adipoq^-/-^* males showed significant increases in the phosphorylation of AMPK, AKT, MAP2K1, and MAPK1 (**Fig. 7A)** and expression of MMP2 and MMP3. No differences were seen for TAGLN or DCN. Aortas of *adipoq^-/-^*females showed a robust upregulation of AMPK phosphorylation and expression of the EGFR, TAGLN, and MMP3 and a reduction in DCN (**Fig. 7B**). Thus, the protein expression profile identified in VSMCs *in vitro* by adiponectin deficiency is relevant *in vivo*.

**Figure 7.**
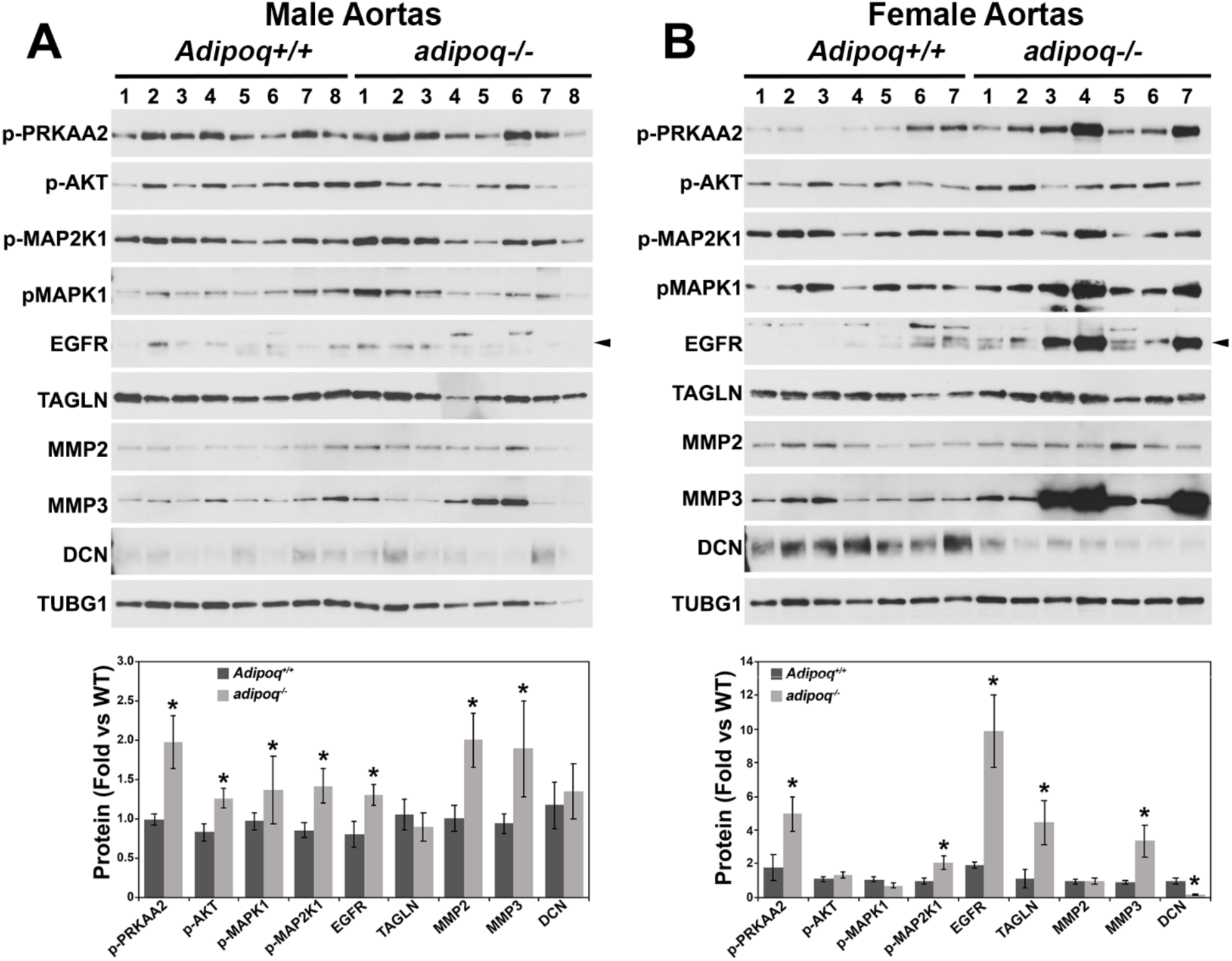
Sex-dependent effects in the expression of cell signaling and ECM markers in aortas of adiponectin deficient mice. Extracts (20-30 μg/well) of aortas of 5-month-old male (A) and female (B) WT and *adipoq^-/-^*mice were separated in 4-20% pre-cast Criterion gels to evaluate changes in protein expression of cell signaling, ECM regulation, and differentiation molecules. * denotes p < 0.05, compared to *Adipoq^+/+^* controls.

## Discussion

Although adiponectin has many protective effects in the cardiovascular system, there have been discrepancies in the literature concerning its effect on atherosclerosis. Nawrocki et al. reported that adiponectin deficiency had no effect on atherosclerosis in male and female *ldlr^-/-^adipoq^-/-^* mice (12). This study also used male *apoe^-/-^adipoq^-/-^* mice that showed less plaque compared with *apoe^-/-^* mice, although differences were not statistically significant. Lindgren et al., who knocked out AR2 together with ApoE, reported that lack of AR2 reduced plaque accumulation in the brachiocephalic artery, but not in the arch or descending aorta of male mice (32). Fujishima et al. showed that deficiency of adiponectin or T-cadherin in *apoe^-/-^*mice promoted plaque accumulation in the descending aorta(12). Adiponectin supplementation in *apoe^-/-^* mice, on the other hand, showed many protective effects including inhibition of lesion formation in the aortic sinus (11, 33, 34), reduced inflammation caused by NF-κB (33), increased superoxide dismutase (SOD) activity, and decreased cholesterol and triglycerides (34). These studies were performed in young mice on HFD; however, the role of adiponectin deficiency during aging is unknown.

In contrast to previous literature, our study revealed novel sex– and age-dependent effects of adiponectin deficiency (homozygote and heterozygote) in atherosclerosis in *apoe^-/-^* mice. HFD reduced plaque in young males and increase it in young females. However, in older mice maintained on a standard chow diet, both sexes showed increased plaque, compared with *apoe^-/-^* controls. Our findings in female aortas and with partial adiponectin deficiency are particularly important because there has only been one other study done measuring plaque in female *apoe^-/-^adipoq^-/-^* mice(3) and no studies done in female or male *apoe^-/-^adipoq^+/-^* mice. Although our study was consistent with Lindgren et al.’s report showing that lack of adiponectin signaling protects against atherosclerosis, this study differs because we show differences in plaque in the arch and descending aorta rather than the brachiocephalic artery. Since only males were investigated by Lindgren et al., it is unknown whether the lack of AR2 would also increase plaque in females. Several other differences in study design between these reports could also explain the differential outcomes. For example, Nawrocki et al. treated 8-week-old animals for 12 weeks with Paigen’s atherogenic diet (35% kcal from fat, 45% kcal from carbohydrates and 1.25% CHO and 0.5% cholic acid), Lindgren et al. treated 8-week-old animals for 14 weeks with lard diet (21% swine lard and 0.15% CHO), and Fujishima et al. treated 8-week-old animals for 12 weeks with HFD (20% fat and 0.15% CHO)(35). However, in the present study, we fed animals with Paigen’s atherogenic diet for only 5 weeks. It is likely that the short treatment time of our study allowed male mice to mount a compensatory mechanism that was not observed in females.

Interestingly, Fujishima et al. showed that T-cadherin mediates the binding of adiponectin to the vascular wall, as the lack of T-cadherin reduced adiponectin expression in the vasculature increasing the levels of this adipokine in circulation(35). T-cadherin has two ligands —adiponectin and LDL—, that have opposite effects in atherosclerosis. T-cadherin binds the high molecular weight form of adiponectin, which is the most active(36). Low levels of adiponectin increase the availability of T-cadherin binding to LDL, leading to pro-atherogenic effects. On the other hand, high levels of circulating adiponectin are not always beneficial and may instead reflect lower binding of the hormone to target organs leading to impaired insulin sensitivity. This phenomenon is known as the “adiponectin paradox”(37). Our data suggest that sex is also a contributing factor to this paradox, as the lack of adiponectin was protective in young males, but not in females.

The effect of adiponectin deficiency on plaque was independent of circulating cholesterol but correlated with a reduction in MCP1, TNF-α, IL-4, CXCL10 in males and an increase in MCP1, IL-6, and CXCL1 in females. These cytokines have major roles in plaque formation; for example, MCP1 mediates the recruitment of monocytes to injury sites in the vasculature and was shown to promote plaque instability in humans(38). In mice, genetic ablation of MCP1 protects from atherosclerosis(39). IL-4 induces MCP1 expression through a redox-dependent mechanism(40), suggesting that reduced IL-4 could be the cause of MCP1 downregulation in males. CXCL10, on the other hand, promotes atherosclerosis by regulating the ratio of effector and regulatory T cells in plaque(40). CSF3, a growth factor involved in the production of mature neutrophils was upregulated in males and reduced in females by adiponectin deficiency. A meta-analysis suggests that CSF3 administration reduces atherosclerosis likely by mobilizing progenitor cells(41).

Similar to previous reports (3, 42), *adipoq^-/-^* cells underwent a switch to the synthetic phenotype with increased proliferation and migration and reduced expression of differentiation markers. This phenotype was associated with heightened expression of glycolytic enzymes at the RNA and protein levels, and impaired OXPHOS. Upregulation of LDH suggests that the conversion of pyruvate into lactate provides a source of NAD needed to maintain a higher glycolytic rate. Analysis of pyruvate metabolism from the RNA sequence data suggests that the activity of the pyruvate dehydrogenase (PDH) complex should be upregulated in *adipoq^-/-^* cells since expression of the pyruvate dehydrogenase kinase (PDK2 and 4), an inhibitor of the PDH complex, is downregulated, while the pyruvate dehydrogenase phosphatase (PDP2), an activator of the PDH complex is upregulated. This suggests that Acetyl-CoA, a central regulator of epigenetic modifications, should be increased. Thus, it is possible that a larger Acetyl-CoA pool in adiponectin deficient cells promotes the upregulation of glycolytic enzymes as well as other genes increased in these cells. Measurements of nicotinamide adenine dinucleotide (NAD) and Acetyl-CoA should be evaluated in future research to elucidate the role of adiponectin in epigenetic modifications in VSMCs.

RNA sequence data of male *adipoq^-/-^* VSMCs also provide insight into possible protective mechanisms of adiponectin deficiency in males. These data also furthered our understanding of the importance of adiponectin in cellular homeostasis while indicating potential key pathways involved in the regulation of the phenotypic switch of VSMCs. We focused on the PI3K/AKT and ECM pathways to identify molecular mechanisms regulating the phenotypic switch and the expression of anti-atherogenic molecules.

Our *in vitro* data show that there are significant differences in the secretory profile, proliferation, migration, and differentiation with the loss of adiponectin. The signaling cascade through AKT, TGF-β, and AMPK are major potential targets for reversing the synthetic phenotype of *adipoq^-/-^*VSMCs. Our data are consistent with a model in which adiponectin activates AMPK, leading to reduced AKT and MAPK1 activation. Through the inhibition of AKT, adiponectin reduces proliferation and migration of VSMCs, and through the inhibition of MAPK1, adiponectin maintains the differentiated state of VSMCs. In *adipoq^-/-^* VSMCs, TGF-β promoted proliferation and migration through AKT activation. Our data agrees with the current literature which shows that TGF-β promotes AKT signaling and that is, as previously mentioned, a cause for increased VSMC proliferation (43). Thus, it is likely that in adiponectin deficient cells, inhibiting TGF-β reduces proliferation by an AKT-dependent mechanism. Additionally, we found that activating AMPK with AdipoRon produced similar effects to AKT and TGF-β inhibition. This is a significant finding because it agrees with previous reports on AdipoRon (44). Also, in agreement with our data, Zhou et al. have shown that globular adiponectin inhibits osteoblastic differentiation of VSMCs through the PI3K/AKT and Wnt/β-catenin pathways(44). This points back to the dysregulation of AKT signaling as the likely primary cause behind the increased proliferation and migration of *adipoq^-/-^* VSMCs which we have shown in this paper.

Our data agrees with work by Ding et al., which showed that adiponectin influences differentiation in human coronary artery smooth muscle cells(3). Initial analysis of differentiation markers in *adipoq^-/-^*cells showed a significant reduction in key proteins such as MHY11, ACTA2, TAGLN, and MYL. AKT induces MYOCD transcription by phosphorylation and inhibition of Foxo3a, a negative regulator of MYOCD transcription (45). Thus, increased AKT activity may explain why MYOCD expression is upregulated. MYOCD and SRF upregulation could also be a compensatory mechanism to correct for the reduced expression of differentiation markers. The reduced effect of MYOCD and SRF suggest that promoter regions of differentiation genes are not accessible (closed chromatin) and/or that these transcription regulators are in the wrong compartment (cytosol instead of nucleus). In fact, we observed a reduction in SRF nuclear localization in adiponectin deficient cells that may explain, in part, the reduced expression of differentiation markers. The role of adiponectin in epigenetic modifications needs further research.

Inhibition of the activity of AKT by TCB and MAPK1 by PD induced a small, but significant increase in ACTA2 expression. However, inhibition of these kinases via activation of adiponectin signaling with AdipoRon and metformin showed no effect on ACTA2, but reduced proliferation and migration of *adipoq^-/-^* cells. Fairaq et al. demonstrated that AdipoRon reduced proliferation in a mTOR dependent mechanism (44). It is possible that the effect of AKT in differentiation is isoform specific. In fact, AKT1 and AKT2 have opposite effects on proliferation and differentiation. Knockdown of AKT1 induces a contractile phenotype, while knockdown of AKT2 a dedifferentiated state(46). TGF-β is known to upregulate the phosphorylation of AKT2 to stimulate differentiation (3, 42). While TCB inhibits all AKT isoforms, it is unknown which AKT isoform is inhibited by the adiponectin/AMPK pathway. It is also possible that the phenotypic switch has more than one arm of regulation. Proliferation may be most impacted by AKT1, while differentiation may be most sensitive to a combination of both AKT2 and MAPK1. This is in line with the current literature, which has indicated that differentiation is regulated by MAPK (47). The MAPK pathway regulates differentiation by phosphorylating (inhibiting) the MYOCD transactivation domain, which leads to reduction in proteins like TAGLN and ACTA2 (48). Also, our finding that AKT is implicated in differentiation, albeit not as strongly as MAPK, agrees with the current literature (39).

With respect to the ECM, molecules regulating TGF-β function were upregulated by adiponectin deficiency, including DCN, MMP14, and TFRC. In fact, the TGF-β pathway was more active in *adipoq^-/-^* cells (increased AKT and SMAD2 phosphorylation), inducing a stronger upregulation of TFRC with TGF-β treatment. However, the upregulation of ACTA and MMP14 in response to TGF-β treatment in WT cells was not observed in *adipoq^-/-^*cells, suggesting that some arms of the TGF-β signaling pathway are impaired. Our data corroborate other studies which also determined that inhibition of TGF-β is detrimental to differentiation (41).

MMP2 and MMP9 were reduced, and MMP3 was upregulated *in vitro*. Our data agree with some findings showing adiponectin reducing MMP9(49), but contradict the results of studies by Wanninger et al. which reported that adiponectin increases MMP9 activity. Notably, however, these studies were conducted in hepatocytes (50). Consistent with our *in vitro* work, the phosphorylation of AKT, MAPK1 and MAP2K1, as well as the expression of MMP2 and MMP3 were upregulated in the aorta of *adipoq^-/-^* male mice, compared to WT. The upregulation of MMP2 and MMP3 is particularly interesting because of its known role in advancing atherosclerotic plaque development (51). Only MPK2K1 phosphorylation and EGFR and MMP3 expression were elevated in females by adiponectin deficiency. The major sex-dependent differences *in vivo* were for MMP3 and DCN, which showed a stronger upregulation in females, compared with males. Unexpectedly, AMPK activity was upregulated in both sexes by adiponectin deficiency, which may signify a compensatory mechanism to curved down plaque progression. It is likely that the robust upregulation of MMP3 and downregulation of DCN impaired this compensatory mechanism in females. The sex-dependent effects of MMP3 and DCN in plaque requires further elucidation in future research.

Overall, we identified protective pathways that may explain the reduced plaque accumulation in young males with adiponectin deficiency. *In vivo*, these mechanisms include the increase in AMPK activity, preservation of DCN expression, decreases in inflammatory markers including MCP1, TNF-α, IL-4, and CXCL10 and estradiol upregulations. It is likely that the dysregulation of the ECM (increased MMP3 and reduced DCN) and increases in pro-inflammatory markers like MCP1, IL-6, and CXCL1 mediates the higher plaque accumulation in females with adiponectin deficiency. *In vitro*, we have shown that adiponectin protects the VSMC phenotype by inhibiting AKT-dependent proliferation and migration, MAPK-dependent dedifferentiation, and TGF-β-dependent ECM regulation. In conclusion, in this paper, we provide novel *in vivo* evidence of the sex differences in the response to adiponectin deficiency in atherosclerosis and provide an *in vitro* molecular mechanism that explains the regulation of the phenotypic switch of VSMCs by adiponectin.

Some limitations of our study include 1) only cells derived from male mice were used to generate RNA sequence and the *in vitro* data, 2) metabolites like NAD and Acetyl-CoA should be evaluated in order to identify the role of adiponectin in epigenetics in future research. 3) aortas of *apoe^-/-^adipoq^-/-^* animals on HFD should be used to better correlate the level of plaque with the changes of protein expression in the aorta.

## Methods

### Animal Model

Single knockout mice for Apoe (B6.129P2-Apoe^tm1Unc^/J, Strain # 2052) and adiponectin (B6.129-Adipoq^tm1Chan^/J, Strain # 8195) were purchased from Jackson Laboratory and crossed to generate *apoe^-/-^adipoq^-/-^*and *apoe^-/-^adipoq^+/-^* experimental animals. Three to four-month-old male and female mice (n=8/group) were used for this study. To accelerate the formation of plaque, animals were fed a modified Paigen’s atherogenic purified HFD for 5 weeks. The HFD contains (in % Kcal) 34.9 fat, 45.1 carbohydrates and 20.1 protein in addition to 0.5% cholic acid. Food and water consumption and body weights were recorded weekly. Upon completion of 5 weeks, animals were euthanized with CO_2_. Blood was collected from the left ventricle and serum stored at –80°C.

### Aortic Plaque Measurement

Aortas of mice fed HFD were isolated, cleaned of periadventitial fat, fixed in 0.2% glutaraldehyde and opened longitudinally for plaque analysis, as reported (52). Aortas were then pinned in a petri dish containing black wax and photographed for the quantification of plaque. This measurement was done by determining the area of plaque, compared to total area of the aorta using ImageJ software. Percent plaque was calculated for the arch and descending aorta.

### Proliferation and Cell Migration Assays

To assess proliferation, WT and *adipoq*^-/-^ cells were seeded in 12 well plates with 5000 cells/well and cultured in media with 10% FBS for 48 h. Cells were then incubated in media with 0.2% FBS for 24 h to synchronize cell growth cycles. Cells were switched back to the 10% FBS media and counted every day in triplicate with a hemocytometer chamber for 7 d. Cell migration was assessed via scratch-wound analysis. Cells were seeded in 6 well plates with grid lines indicating where scratch-wounds will occur and grown to confluency. A 1 mL pipette tip was used to create the scratch-wound and pictures were taken at the beginning and after 12 and 24 h of treatment. Migration was assessed by measuring scratch-wound width using photoshop and was presented as percent-reduction. The percent closure was calculated as follows: |(cm Xh-cm 0h)*100)/cm 0h)| with 0 representing the open wound at the beginning and 100% a completely close wound. Proliferation and scratch wound analyses were done on WT and *adipoq^-/-^* cells with no treatment and *adipoq^-/-^*cells treated with recombinant adiponectin (10 μg/mL) metformin (5 mM), AdipoRon (10 μM), erlotinib (Erl, epidermal growth factor receptor – EGFR inhibitor, 100 nM), triciribine (TCB, AKT inhibitor, 1 μM), PD-98059 (PD, mitogen-activated protein kinase kinase MEK/MAP2K1 inhibitor, 10 μM) and SB-252334 (SB, TGF-β receptor inhibitor, 5 μM).

### Statistics

Western blot results were analyzed using ImageJ. Protein expression was determined in at least 3 independent experiments, adjusted by the level of loading control (tubulin, GAPDH, or β-actin) and data were analyzed by comparing WT and *adipoq^-/-^* cells (2 groups) using two-way t-tests in JMPPro15 (version 15.0.0). For cell culture experiments utilizing time as a variable (proliferation curve and scratch wound assay) a mixed model repeated measures test (genotype, time, and genotype* time) was used. JMPPro15 was used for ANOVA and Tukey’s Post Hoc analysis for comparison of body weight, food intake, and water consumption for in vivo studies. For box and whisker plots, some individual values are similar so that only one dot appears on the graphs. Thus, data points on graphs are not indicative of sample size. Values are given as mean ± standard deviation of mean (SD), except for body weight, food and water which are given as mean ± standard error (SE) and p < 0.05 is considered statistically significant.

### Study Approval

Experiments using mice were performed in compliance with the Public Health Service Policy and approved by the Animal Care and Use Committee at Florida State University. Protocol # 201900033.

### Author Contributions

The authors’ contributions were as follows—GS conceived the study and obtained funding; AEC, AMC and VU conducted the in vivo experiment; GS, AEC, AMC, AI and RD conducted the *in vitro* experiments; GS, AEC, AMC, AI, PK and JD were involved in data analysis; GS, AEC, AMC were involved in manuscript writing; GS had primary responsibility for final content; all authors reviewed and approved the final manuscript.

## Acknowledgements

This research was supported by the US Department of Agriculture (USDA-AFRI, GRANT12444832) and the Florida Department of Health, James and Esther King Biomedical Research Program (9JK01).

## Major Resources Table

In order to allow validation and replication of experiments, all essential research materials listed in the Methods should be included in the Major Resources Table below. Authors are encouraged to use public repositories for protocols, data, code, and other materials and provide persistent identifiers and/or links to repositories when available. Authors may add or delete rows as needed.

### Animals (in vivo studies)

| Species | Vendor or Source | Background Strain | Sex | Persistent ID / URL |
| --- | --- | --- | --- | --- |
| wild-type mouse | The Jackson Laboratory | C57BL/6J strain | M, F | Stock#000664; RRID:MGI:5650797 |
| Adipo<sup>tm1Chan</sup> mouse | The Jackson Laboratory | C57BL/6J background | M, F | Stock#008195; RRID:IMSR_JAX:008195 |
| ApoE<sup>-/-</sup> mouse | The Jackson Laboratory | C57BL/6J background | M, F | Stock#002052; RRID:IMSR_TAC:APOE |
| ApoE<sup>-/-</sup>Adipoq<sup>-/-</sup> | Salazar Lab | C57BL/6J background | M, F | Generated in the Salazar's lab |

### Antibodies

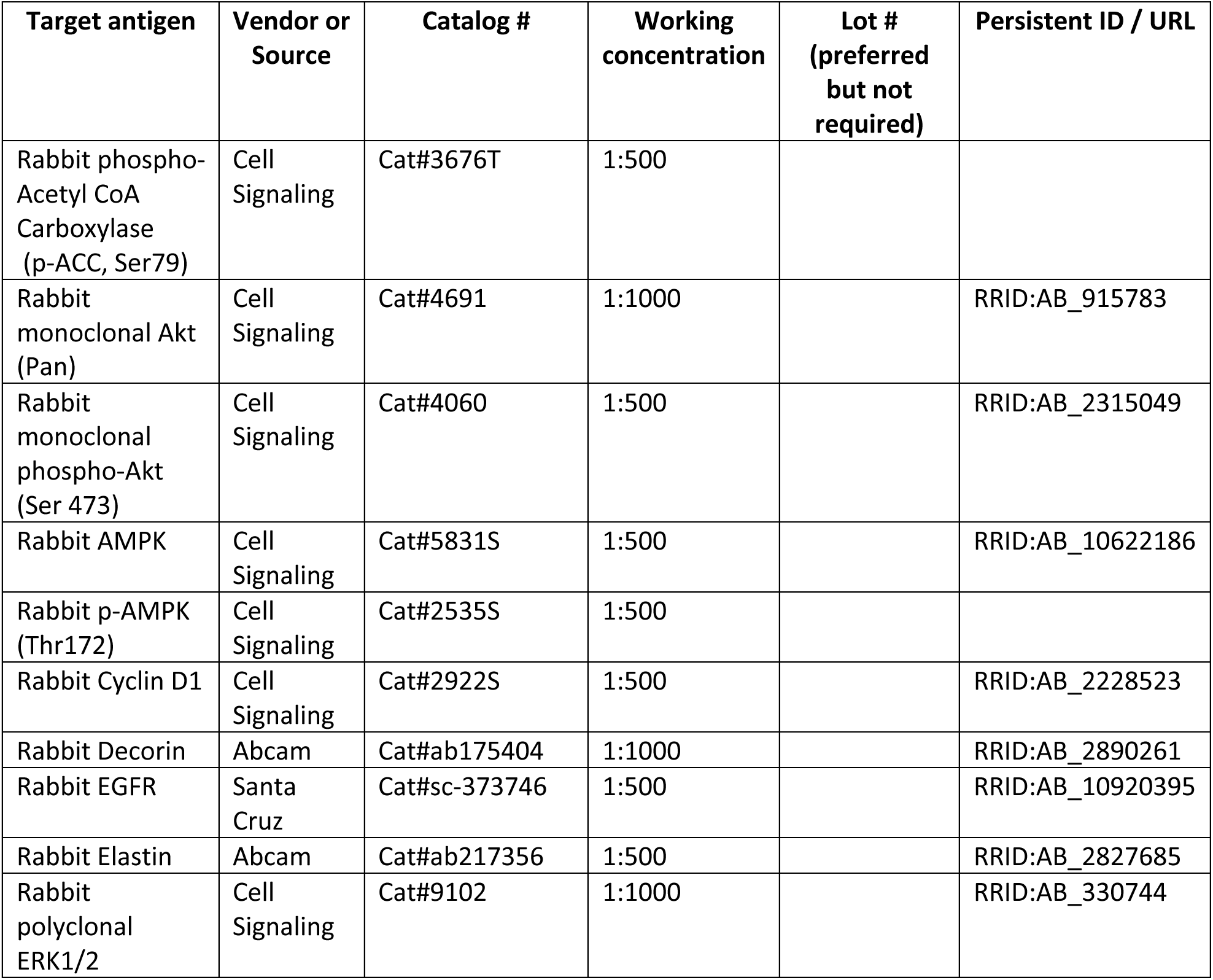
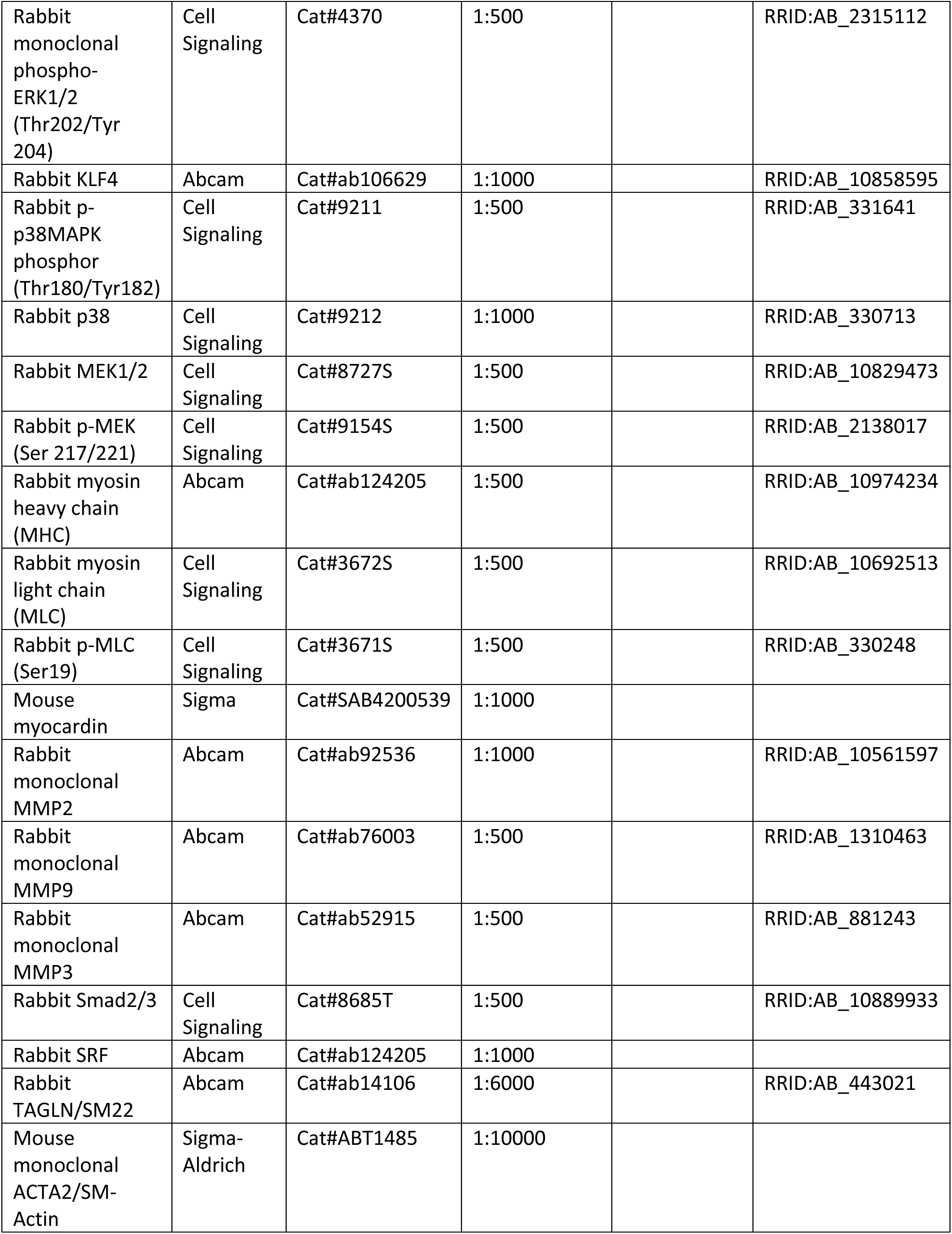
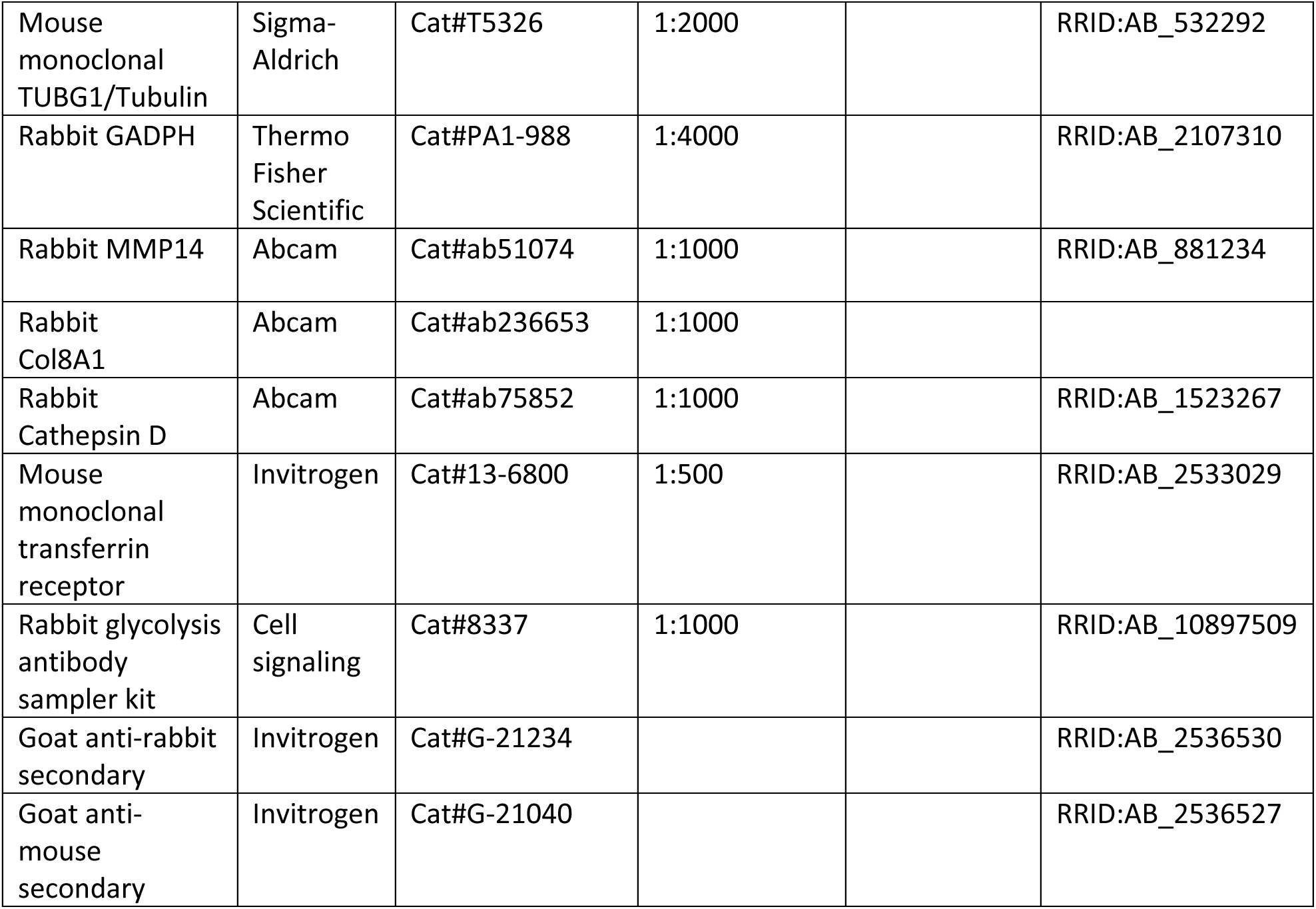

### Cultured Cells

| ***Name*** | ***Vendor or Source*** | ***Sex (F, M, or unknown)*** | ***Persistent ID / URL*** |
| --- | --- | --- | --- |
| Mouse aortic C57BL/6J VSMCs (MASMs) | Salazar's Lab | Male |  |
| Mouse aortic *adipoq*<sup>-/-</sup> VSMCs (MASMs) | Salazar's Lab | Male |  |

### Other

| ***Description*** | ***Source / Repository*** | ***Persistent ID / URL*** |
| --- | --- | --- |
| Adiponectin (recombinant, mouse) | BioVendor | Cat#101215-406 |
| AdipoRon | Sigma Aldrich | Cat# SML0998-5MG |
| TCB (Triciribine, Akt inhibitor V) | Millipore | Cat#124012-1MG |
| 2',7'-dichlorodihydrofluorescein diacetate (H<sub>2</sub>DCFDA) | Invitrogen | Cat#C6827 |
| DHE | Life Technologies | Cat#D11347 |
| MitoSox | Invitrogen | Cat#M36008 |
| Erlotinib | Sigma Aldrich | Cat#SML2156 |
| EZ Link Sulfo-NHS-Biotin | Pierce | Cat#21217 |
| Metformin Hydrochloride | Sigma Aldrich | Cat#PHR1084-500MG |
| Immobilized Neutravidin | Pierce | Cat#29200 |
| PD-98059 MEK inhibitor | Enzo Life Sciences | Cat#BML-E1360-0005 |
| SB-525334 TGF-β inhibitor | VWR | Cat#103539118 |
| Silver Stain Plus kit | Biorad | Cat#161-0449 |
| Lipid Plus Piccolo kit | McKesson | Cat#07P0205 |
| Milliplex Mouse Cytokine/Chemokine Magnetic Kit | Millipore | Cat#MCYTMAG-70K-PX32 |
| Milliplex Mouse CVD Magnetic Bead Panel 1 | Millipore | Cat#MCVD1MAG-77K |
| Milliplex Multi-species Hormone Magnetic Bead Panel | Millipore | Cat#MSHMAG2-21K |
| Milliplex Mouse MMP Magnetic Bead Panel 1 | Millipore | Cat#MMMP1MAG-79K |
| ImageJ 6.0 | Schneider et al., 2012 | https://imagej.nih.gov/ij/ |
| xPONENT |  | https://www.luminexcorp.com/xponent/ |
| IBM (International Business Machines) SPSS 25 (Statistical Package for the Social Sciences version 25) |  | https://www.ibm.com/docs/en/spss-statistics/25.0.0 |

