## Supplementary data for "The Duality of Adiponectin and the Role of Sex in Atherosclerosis"

### Supplemental Methods and Figures

*Genotyping*

Mice were gently sedated with isoflurane to have tail tip collected and ears punched for identification. DNA from tails was isolated using the Quanta Accustart II Mouse Genotyping Kit (Quanta Biosciences, 95091). PCR was then run in a Mastercycler Nexus Gradient (Eppendorf). Primers sequences were obtained from Jackson Laboratory genotyping protocols database and were purchased from Sigma Aldrich. The sequence of the primers is shown in Table S1.

**Table S1. Sequence of genotyping primers**

| Gene | **Primer ID** | **Sequence** | **Primer type** |
|---|---|---|---|
| ApoE | oIMR0180 | GCC TAG CCG AGG GAG AGC CG | Common Forward |
| | oIMR0181 | TGT GAC TTG GGA GCT CTG CAG C | WT reverse |
| | oIMR0182 | GCC GCC CCG ACT GCA TCT | Mutant reverse |
| Adipoq | 28949 | GGT GGC TCA CAA CCA TTC A | Common Forward |
| | 28950 | CTC CCA GGA GGT CTT CAT CA | WT reverse |
| | 24955 | CTT CCT GAC TAG GGG AGG AG | Mutant reverse |

*Preparation of Mouse VSMCs*

VSMCs were harvested from aortas of male C57Bl/6 WT and *adipoq*$^{-/-}$ mice using an explant method, as reported [1]. Aortas were isolated and treated with collagenase (150 U/ml, Worthington, 4202) for 30 min at 37°C to remove the tunica adventitia. The aorta was then cut lengthwise, and the intima was removed with a sterile cotton swab. The remaining media layer was cut into small pieces and allowed to attach to the culture plate for at least 3 days. Explants were cultured until plates were 80% confluent. Cells were expanded, frozen using 10% DMSO (Sigma-Aldrich, 1001731596) and 90% FBS (Seradigm, 97068-075) and stored in liquid nitrogen.

*Cell culture*

Cells were grown in T75 cell culture flasks (Greiner; 658170) and cultured until passage 12 in Dulbecco's Modified Eagle Medium (DMEM) (Corning; 10-014-CV) supplemented with 10% FBS, and glutamine (2 mM), penicillin (100 U/L), and streptomycin (100 mg/ml) (Corning; 30-009-CI). Cells were cultured at 37°C in a 5% $CO_2$ incubator, and the medium was changed every other day. For treatment, cells were starved in DMEM with 0.2% FBS for 24 h before subsequent treatments.

*Cell Lysis and Sample Preparation*

Plates were placed on ice and washed twice with cold PBS containing calcium and magnesium (PBS Ca/Mg) (1 mM $MgCl_2$, 0.1 mM $CaCl_2$) and were lysed with 120 µl of Buffer A (50 mM HEPES, pH 7.4, 150 mM NaCl, 1 mM EGTA and 0.1 mM $MgCl_2$) containing 1% Triton X-100 (Sigma-Aldrich, X100-500ML), 2 mM sodium orthovanadate (Enzo Life Sciences, 400-032-G025), 10 mM sodium pyrophosphate (Sigma-Aldrich, 221368), 10 mM sodium fluoride (J.T. Baker, 368-01), and protease inhibitor cocktail (Sigma, P340-5ML). Samples were sonicated 3 times for 10 s using a QSonica sonicator and protein analysis was completed with Bradford reagent (BioRad, 50000006) in a spectrophotometer. Samples were adjusted by protein content (20-50 µg) then separated in either 4-20% acrylamide precast Criterion gels (BioRad, 5671094) or homemade 8%, 10%, or 12% SDS-PAGE gels.

*Western Blot*

Gels were transferred to PVDF membrane (Thermo Scientific, PI88518) using the Thermo Scientific Owl HEP-1 transfer system and membranes blocked for 20 min in TBS (150 mM NaCl, 2 mM KCl, 25 mM Tris, pH 7.4) containing 1.5% non-fat dry milk (Bio-Rad, 170-6406). Membranes were washed in TBS and incubated with primary antibody from 1-2 h to overnight. Membranes were then washed 3 times, 10 min each, with TBS-T (TBS plus 0.05% T-X100) and incubated with secondary antibodies in block buffer for 45 min. After 3 washes in TBS-T, membranes were incubated with Pierce ECL Western Blotting Substrate (Thermo Fisher).

*Biotinylation*

Cells were starved for 24 h in 0.2% FBS and then were washed twice with PBS Ca/Mg and incubated with 0.5 mg/ml sulfo-NHS-biotin in PBS Ca/Mg for 30 min on ice [1, 2]. After two washes with cold PBS-CM, cells were incubated in PBS Ca/Mg containing 50 mM $NH_4Cl$ for 10 min to stop the biotinylation reaction. Cells were washed 3 times with PBS Ca/Mg and lysed with 500 μl lysis buffer for 30 min on ice. Cell extracts were transferred to 1.7 ml microcentrifuge tubes and centrifuged for 10 min at 22,000 x g. Supernatant was collected into new 1.7 ml tubes and incubated with 30 μl NeutrAvidin™ beads for 2 h at 4ºC in a rotating shaker. Beads were washed 5 times for 5 min each, resuspended in 30 μl loading buffer and separated in 4-20% precast gels.

*Zymography*

For zymography measuring gelatinase activity, 8% SDS-PAGE gel was made with the addition of 1% gelatin (Sigma, G7041-100G). Cells were treated in plain media, which was concentrated with Amicon Ultra-0.5 column with a 10 kDa molecular weight cutoff. Samples of concentrated media were separated under non-reducing conditions and without heating to preserve the structure of the enzymes. Upon completion, the gel was washed in 2.5% Triton X-100 in water for 30 min, with one change of the solution after 15 min. The gels were then rinsed in substrate buffer (500 mM Tris-HCl, pH 7.8, 2 mM NaCl, 50 mM $CaCl_2$) overnight and incubated in staining buffer (0.25% Coomassie blue R-250, 5% methanol and 10% acetic acid) for 2 h. After the gel was fully dyed, it was rinsed in destaining buffer (10% methanol, 5% acetic acid) overnight.

*Oroboros Oxygraph System*

To determine the function of the mitochondrial complexes associated with oxidative phosphorylation we used the Oroboros Oxygraph System (Oroboros Instruments, Innsbruck Austria), as previously reported [3]. WT and *adipoq*$^{-/-}$ cells were grown to confluency, and approximately 1-2 million cells were resuspended in mitochondrial respiration medium MiR05 (0.5 mM EGTA, 3 mM $MgCl_2$, 60 mM lactobionic acid, 20 mM taurine, 10 mM $KH_2PO_4$, 20 mM HEPES, 110 mM D-sucrose and 1g/L fatty acid free BSA, pH 7.1), supplemented with 20 mM creatine monohydrate, and added to each chamber.

Digitonin (4.05 µM) was used to permeabilize the plasma membranes. To stimulate electron flow through Complex 1, malate (2 mM) and glutamate (10 mM) were added to the chambers to produce NADH (state 2 respiration). To measure ADP-stimulated oxygen consumption by Complex 1 (state 3 respiration), ADP was added (4 mM) to the chambers. Next, succinate (10 mM) was added to stimulate electron flow through Complex II as well, followed by rotenone (10 µM) to inhibit Complex I, thus measuring Complex II respiration independent of Complex I. Finally, complex IV respiration was assessed using N, N, N', N'-Tetramethyl-p-phenylenediamine dihydrochloride (TMPD, 0.4 mM), 2 mM ascorbate to prevent TMPD auto-oxidation, and 5 µM antimycin A to inhibit electron flow through complex III. Cytochrome c (10 µM) was added to test mitochondrial membrane integrity, and any samples with a relevant cytochrome c release were excluded. The respiration rate was expressed as pmol oxygen consumed per second and normalized to protein content in the O2k-Chamber (pmols/s/mg protein), measured using a Pierce BCA Protein Assay (Thermo Fisher Scientific).

*Magnetic bead assay*

A magnetic bead kit, Mouse Cytokine/Chemokine Panel (MCYTOMAG-70K), was obtained from Sigma-Aldrich to assess inflammatory blood markers. Plasma samples from male and female *apoe*$^{-/-}$ and *apoe*$^{-/-}$*adipoq*$^{-/-}$ mice (8 per group per sex) were used for this assay. Samples were diluted (1:1 with provided Assay Buffer) before addition to the provided 96-well plate. The plate was washed with wash buffer and incubated on a shaker at room temperature (20-25ºC) for 10 m. The solution was decanted and standards (25 µl/well) or controls (25 µl/well) added to appropriate wells, then assay buffer (25 µl/well), matrix solution (25 µl/well), and diluted samples (25 µl/well) to sample wells, and lastly the magnetic beads (25 µl/well) were added. Plates were incubated 16-20 h at 4ºC. The following day, well contents were removed and 2 washes with 200 µl/well wash buffer were performed. Detection antibodies (25 µl/well) were added, and plate was incubated covered with aluminum foil on a shaker for 1 h. Streptavidin-Phycoerythrin (25 µl/well) was added, and plate was again covered with foil for a 30 m incubation with gentle agitation. Contents were removed and washed twice with wash buffer (200 µl/well).

Sheath/drive fluid was added to each well (150 µl) and 96 well plate was read using a MAGPIX system (Luminex Corporation).

*Nuclear Fractionation*

After starvation for 24 h, cells were washed twice with PBS Ca/Mg and resuspended in 300 µl of hypotonic buffer (20 mM Tris, pH 7.4, 3 mM $MgCl_2$, 10 mM NaCl plus 10 µl/ml protease inhibitor cocktail) for nuclear fractionation, as previously reported [4]. Extracts were collected using a cell scraper and transferred to pre-chilled 1.7 ml microcentrifuge tubes and incubated on ice for 15 min. Then 15 µl of 10% NP40 was added to the samples followed by vortex for 10 s. The samples were centrifuged at 4ºC 22,000 x g for 10 min. The supernatant was saved into pre-chilled 1.7 ml microcentrifuge tubes labeled Cytosolic Fraction. The pellet was washed with 500 µl hypotonic buffer and the excess buffer was removed. The nuclear fraction samples were centrifuged for 1 min and excess buffer was removed. The pellet was suspended in 80 µl of nuclear extraction buffer (100 mM Tris, pH 7.4, 100 mM NaCl, 1 mM EDTA, 10% glycerol, 1 mM EGTA, 0.1% SDS. 0.5% deoxycholate, 1% TritonX-100, and 10 µl/ml protease inhibitor cocktail) and incubated on ice for 30 min, vortexing every 10 min. Finally, the pellet samples were centrifuged for 20 min at 22,000 x g at 4˚C. The supernatant was collected and stored in pre-chilled 1.7 mL microcentrifuge tubes labeled Nuclear Fraction.

*Piccolo Lipid Analysis*

Blood was collected via cardiac puncture and spun down at 5000 x g for 5 minutes to separate the serum. Serum was removed and stored separately in 1.7 mL microcentrifuge tubes until all samples were collected. Due to the fatty nature of the serum from the animals being on high fat diet, the serum was diluted 1:10 in PBS. Once diluted, 100 µl of the serum was added to the Piccolo lipid panel cartridge (McKesson, 07P0205) and read.

### Supplemental Figure Legends

**Figure S1. Body weight, food, and water intake in HFD-treated mice.** Body weight (A and D), water intake (B and E) and food intake (C and F) were measured weekly in male and female *apoe*$^{-/-}$*adipoq*$^{+/+}$, *apoe*$^{-/-}$*adipoq*$^{+/-}$, and *apoe*$^{-/-}$*adipoq*$^{-/-}$ mice over the course of HFD treatment (5 weeks). * denotes $p < 0.05$, compared to *apoe*$^{-/-}$*adipoq*$^{+/+}$, controls.

**Figure S2. Age impairs the protective effect of adiponectin deficiency in male mice.** *apoe*$^{-/-}$*Adipoq*$^{+/+}$, *apoe*$^{-/-}$*adipoq*$^{+/-}$ and *apoe*$^{-/-}$*adipoq*$^{-/-}$ male and female mice were aged to 1 year and aortas were isolated for en face analysis of plaque (A). Plaque area was measured in the arch (B) and descending aorta (C) using ImageJ. * denotes $p < 0.05$, compared to *apoe*$^{-/-}$*adipoq*$^{+/+}$. N=6-7 for males. For females, n=8 for WT females, n=6 for heterozygotes and n=4 for homozygotes.

**Figure S3. RNA sequence analysis of adiponectin deficient cells.** RNA sequence data of male *Adipoq*$^{+/+}$ and *adipoq*$^{-/-}$ VSMCs (n=4 samples per genotype) are presented as a volcano plot of the gene distribution based on the log2 (*adipoq*$^{-/-}$/WT) for RNA expression (A). Log2 (fold change) >1 were upregulated (red) and <1 were downregulated (blue) genes. Changes in cellular processes (B) and DEGs of the most enriched pathways (C) show strong changes in signal transductions, in particular the PI3K/AKT pathway. Molecular interactions of the most altered pathways are shown in D.

**Figure S4. Changes in RNA expression in cell signaling, ECM and glycolytic pathways induced by adiponectin deficiency.** Bar graphs representing log2 (fold change) for signaling (A) differentiation (B), ECM (C), glycolysis (D), MMPs (E) and TGF-$\beta$ (F) pathways.

**Fig. S5. EGFR does not regulate the upregulation of AKT signaling by adiponectin deficiency** *Adipoq*$^{+/+}$ and *adipoq*$^{-/-}$ VSMCs were starved for 24 h with 0.2% FBS and treated with 100 nM EGF for 0-60 m (A) and total cell extracts probed for AKT and MAPK1 phosphorylation (A-C). Biotinylation shows changes in cell surface levels of EGFR, CAV-

1, and TFRC (D-F). * denotes $p < 0.05$, compared to *Adipoq*$^{+/+}$ controls. N= 3 independent experiments.

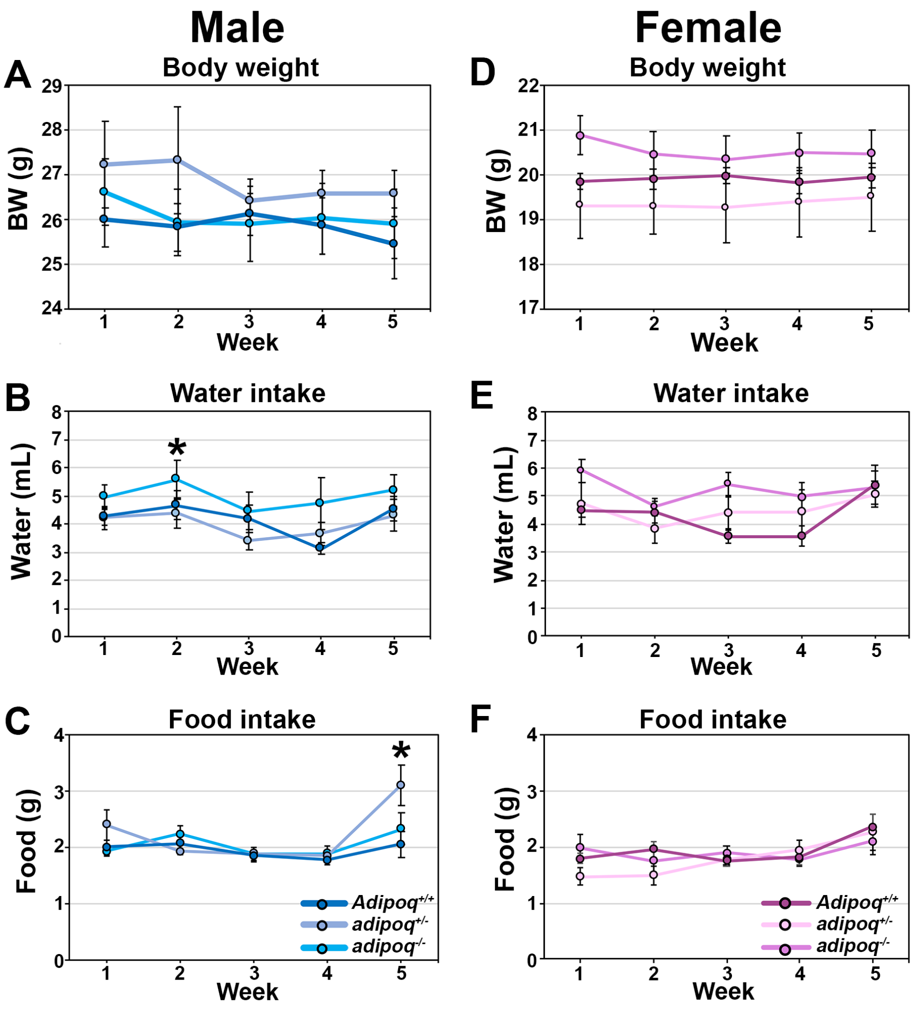

Male
Female
A
Body weight
BW (g)
Week
D
Body weight
BW (g)
Week
B
Water intake
Water (mL)
Week
E
Water intake
Water (mL)
Week
C
Food intake
Food (g)
Week
*Adipoq+/+*
*adipoq+/-*
*adipoq-/-*
F
Food intake
Food (g)
Week
*Adipoq+/+*
*adipoq+/-*
*adipoq-/-*


Figure S1

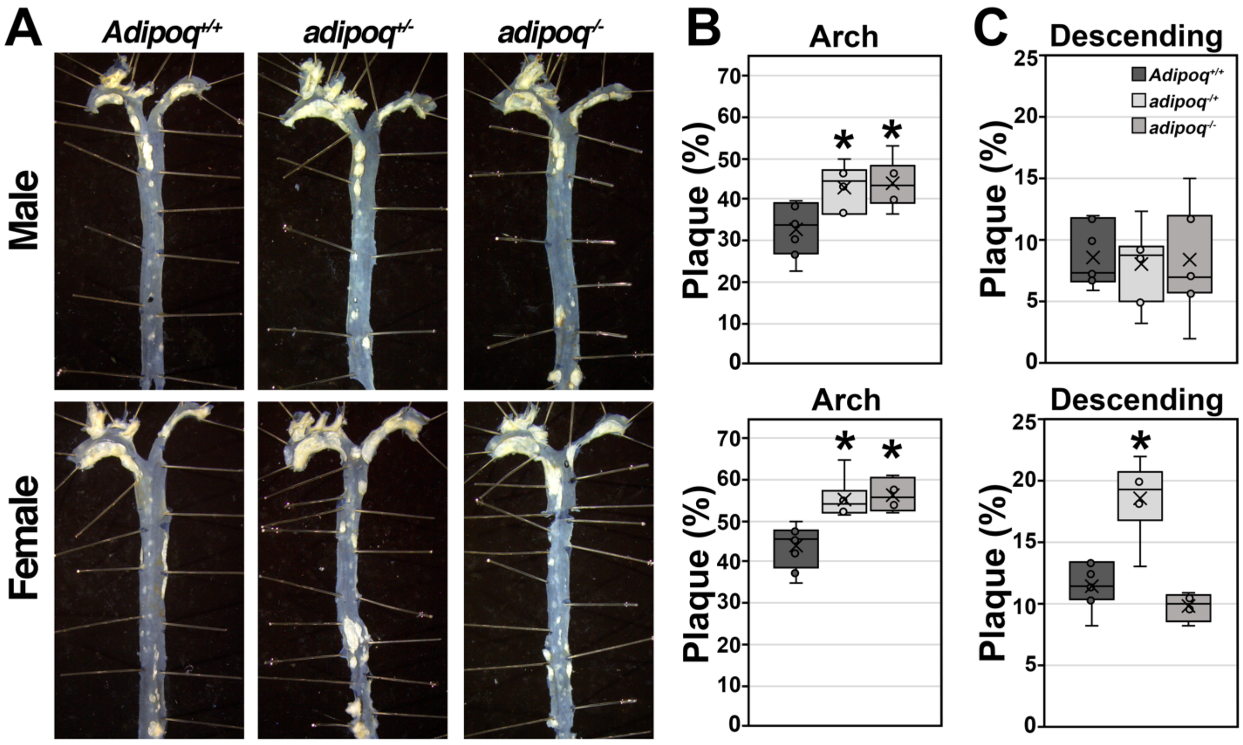

A
Adipoq+/+
adipoq+/-
adipoq-/-
Male
Female
B
Arch
Plaque (%)
Arch
Plaque (%)
C
Descending
Adipoq+/+
adipoq-/+
adipoq-/-
Plaque (%)
Descending
Plaque (%)


Figure S2

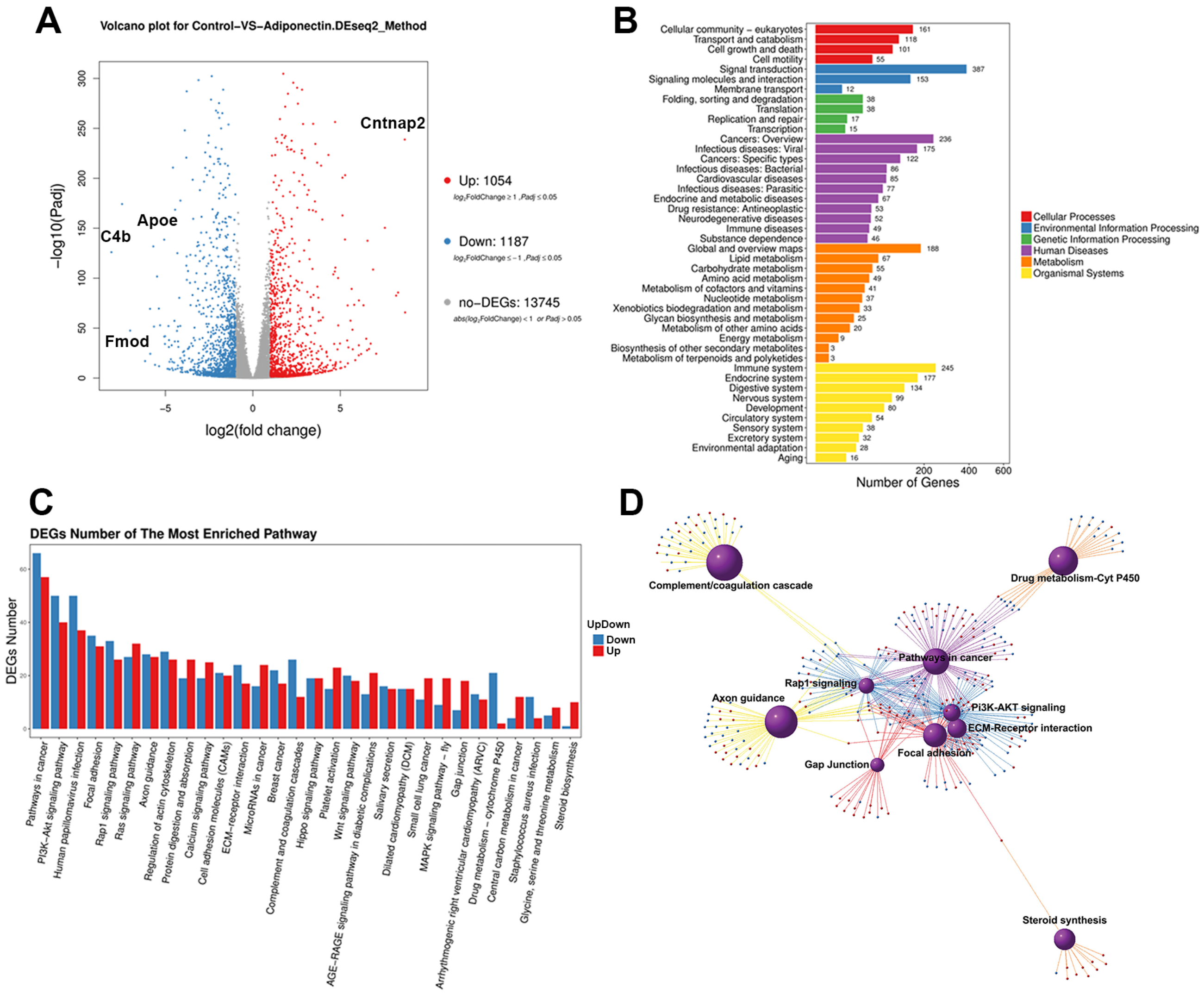
A
Volcano plot for Control-VS-Adiponectin.DEseq2_Method
-log10(Padj)
log2(fold change)
Cntnap2
Apoe
C4b
Fmod
Up: 1054
Down: 1187
no-DEGs: 13745
B
Number of Genes
Cellular community - eukaryotes
Transport and catabolism
Cell growth and death
Cell motility
Signal transduction
Signaling molecules and interaction
Membrane transport
Folding, sorting and degradation
Translation
Replication and repair
Transcription
Cancers: Overview
Infectious diseases: Viral
Cancers: Specific types
Infectious diseases: Bacterial
Cardiovascular diseases
Infectious diseases: Parasitic
Endocrine and metabolic diseases
Drug resistance: Antineoplastic
Neurodegenerative diseases
Immune diseases
Substance dependence
Global and overview maps
Lipid metabolism
Carbohydrate metabolism
Amino acid metabolism
Metabolism of cofactors and vitamins
Nucleotide metabolism
Xenobiotics biodegradation and metabolism
Glycan biosynthesis and metabolism
Metabolism of other amino acids
Energy metabolism
Biosynthesis of other secondary metabolites
Metabolism of terpenoids and polyketides
Immune system
Endocrine system
Digestive system
Nervous system
Development
Circulatory system
Sensory system
Excretory system
Environmental adaptation
Aging
Cellular Processes
Environmental Information Processing
Genetic Information Processing
Human Diseases
Metabolism
Organismal Systems
C
DEGs Number of The Most Enriched Pathway
DEGs Number
UpDown
Down
Up
Pathways in cancer
PI3K-Akt signaling pathway
Human papillomavirus infection
Focal adhesion
Rap1 signaling pathway
Ras signaling pathway
Axon guidance
Regulation of actin cytoskeleton
Protein digestion and absorption
Calcium signaling pathway
Cell adhesion molecules (CAMs)
ECM-receptor interaction
MicroRNAs in cancer
Breast cancer
Complement and coagulation cascades
Hippo signaling pathway
Platelet activation
Wnt signaling pathway
AGE-RAGE signaling pathway in diabetic complications
Salivary secretion
Dilated cardiomyopathy (DCM)
Small cell lung cancer
MAPK signaling pathway - fly
Gap junction
Arrhythmogenic right ventricular cardiomyopathy (ARVC)
Drug metabolism - cytochrome P450
Central carbon metabolism in cancer
Staphylococcus aureus infection
Glycine, serine and threonine metabolism
Steroid biosynthesis
D
Complement/coagulation cascade
Drug metabolism-Cyt P450
Pathways in cancer
Rap1 signaling
Axon guidance
PI3K-AKT signaling
ECM-Receptor interaction
Focal adhesion
Gap Junction
Steroid synthesis

Figure S3

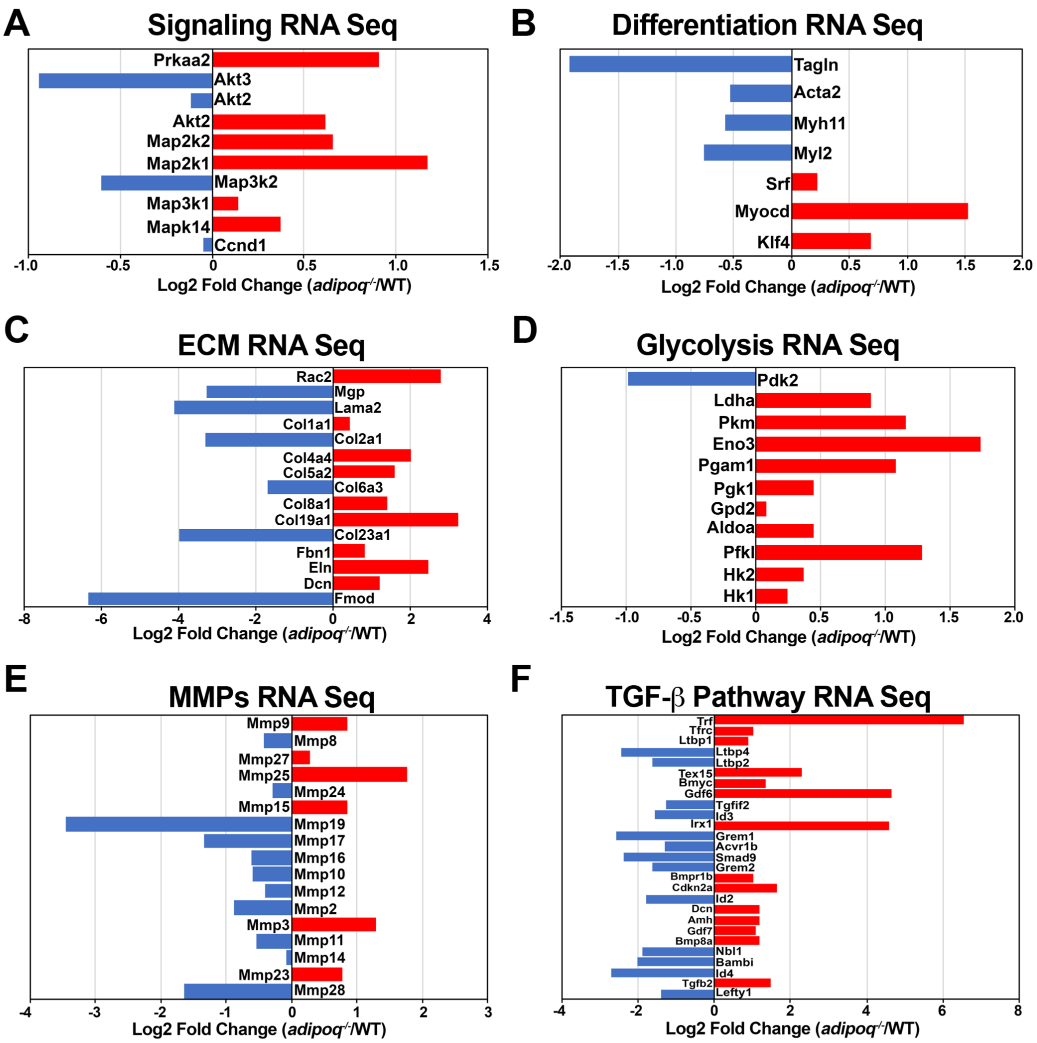
A
Signaling RNA Seq
Prkaa2
Akt3
Akt2
Akt2
Map2k2
Map2k1
Map3k2
Map3k1
Mapk14
Ccnd1
Log2 Fold Change (adipoq-/-/WT)
B
Differentiation RNA Seq
Tagln
Acta2
Myh11
Myl2
Srf
Myocd
Klf4
Log2 Fold Change (adipoq-/-/WT)
C
ECM RNA Seq
Rac2
Mgp
Lama2
Col1a1
Col2a1
Col4a4
Col5a2
Col6a3
Col8a1
Col19a1
Col23a1
Fbn1
Eln
Dcn
Fmod
Log2 Fold Change (adipoq-/-/WT)
D
Glycolysis RNA Seq
Pdk2
Ldha
Pkm
Eno3
Pgam1
Pgk1
Gpd2
Aldoa
Pfkl
Hk2
Hk1
Log2 Fold Change (adipoq-/-/WT)
E
MMPs RNA Seq
Mmp9
Mmp8
Mmp27
Mmp25
Mmp24
Mmp15
Mmp19
Mmp17
Mmp16
Mmp10
Mmp12
Mmp2
Mmp3
Mmp11
Mmp14
Mmp23
Mmp28
Log2 Fold Change (adipoq-/-/WT)
F
TGF-β Pathway RNA Seq
Trf
Tfrc
Ltbp1
Ltbp4
Ltbp2
Tex15
Bmyc
Gdf6
Tgfif2
Id3
Irx1
Grem1
Acvr1b
Smad9
Grem2
Bmpr1b
Cdkn2a
Id2
Dcn
Amh
Gdf7
Bmp8a
Nbl1
Bambi
Id4
Tgfb2
Lefty1
Log2 Fold Change (adipoq-/-/WT)

Figure S4

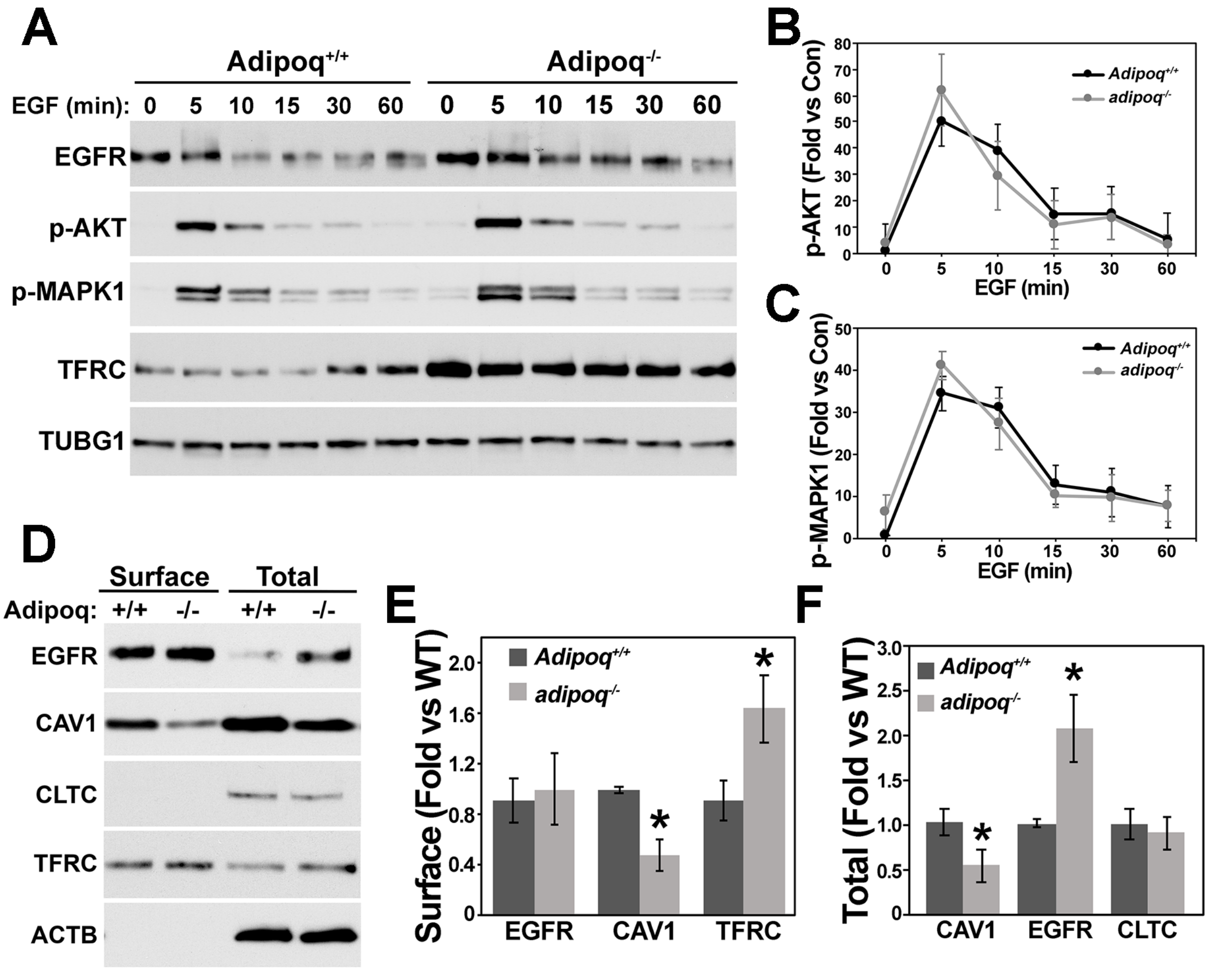
A
Adipoq+/+
Adipoq-/-
EGF (min): 0 5 10 15 30 60 0 5 10 15 30 60
EGFR
p-AKT
p-MAPK1
TFRC
TUBG1
B
p-AKT (Fold vs Con)
EGF (min)
Adipoq+/+
adipoq-/-
C
p-MAPK1 (Fold vs Con)
EGF (min)
Adipoq+/+
adipoq-/-
D
Surface
Total
Adipoq: +/+ -/- +/+ -/-
EGFR
CAV1
CLTC
TFRC
ACTB
E
Surface (Fold vs WT)
Adipoq+/+
adipoq-/-
EGFR CAV1 TFRC
F
Total (Fold vs WT)
Adipoq+/+
adipoq-/-
CAV1 EGFR CLTC

Figure S5
